# Content-aware frame interpolation (CAFI): Deep Learning-based temporal super-resolution for fast bioimaging

**DOI:** 10.1101/2021.11.02.466664

**Authors:** Martin Priessner, David C.A. Gaboriau, Arlo Sheridan, Tchern Lenn, Jonathan R. Chubb, Uri Manor, Ramon Vilar, Romain F. Laine

## Abstract

The development of high-resolution microscopes has made it possible to investigate cellular processes in 4D (3D over time). However, observing fast cellular dynamics remains challenging as a consequence of photobleaching and phototoxicity. These issues become increasingly problematic with the depth of the volume acquired and the speed of the biological events of interest. Here, we report the implementation of two content-aware frame interpolation (CAFI) deep learning networks, Zooming SlowMo (ZS) and Depth-Aware Video Frame Interpolation (DAIN), based on combinations of recurrent neural networks, that are highly suited for accurately predicting images in between image pairs, therefore improving the temporal resolution of image series as a post-acquisition analysis step. We show that CAFI predictions are capable of understanding the motion context of biological structures to perform better than standard interpolation methods. We benchmark CAFI’s performance on six different datasets, obtained from three different microscopy modalities (point-scanning confocal, spinning-disk confocal and confocal brightfield microscopy). We demonstrate its capabilities for single-particle tracking methods applied to the study of lysosome trafficking. CAFI therefore allows for reduced light exposure and phototoxicity on the sample and extends the possibility of long-term live-cell imaging. Both DAIN and ZS as well as the training and testing data are made available for use by the wider community via the ZeroCostDL4Mic platform.

## Introduction

Live-cell imaging is a powerful tool to study dynamic cellular processes by capturing spatio-temporal organization of biological micro-environments. For this, the imaging speed of a microscopy acquisition needs to be sufficiently high to observe cellular processes and dynamic patterns accurately, which sometimes compromises the signal-to-noise ratio (SNR), resolution of the acquired images and/or viability of the sample. Improving the SNR or increasing the dimensionality of the data recording (e.g., 4D (3D+t) acquisitions) provides better context but slows down the recording speed making it more difficult, if not impossible, to capture and understand dynamic processes.

Although classical mathematical interpolation techniques such as bilinear (BIL) or bicubic (BIC) can artificially increase the temporal image density as a post-acquisition step, those methods do not provide more information about the sample dynamics. A smarter interpolation tool would allow for a time-course acquisition to be re-sampled with higher temporal sampling in a fashion that would be content-aware (2) with respect to the dynamics observed and would therefore provide accurate predictions of the missing temporal frames, thereby enabling effectively higher temporal resolution imaging (see Figure 1a). Such approaches would effectively lower the illumination dose on the specimen, reducing phototoxicity and photobleaching and allowing longer recordings without compromising cell health or data quality. Therefore, developing computational tools that increase the temporal image resolution in a content-aware fashion has the potential to be transformative and can push the capabilities of any speed-limited microscopy modality.

**Fig. 1.**
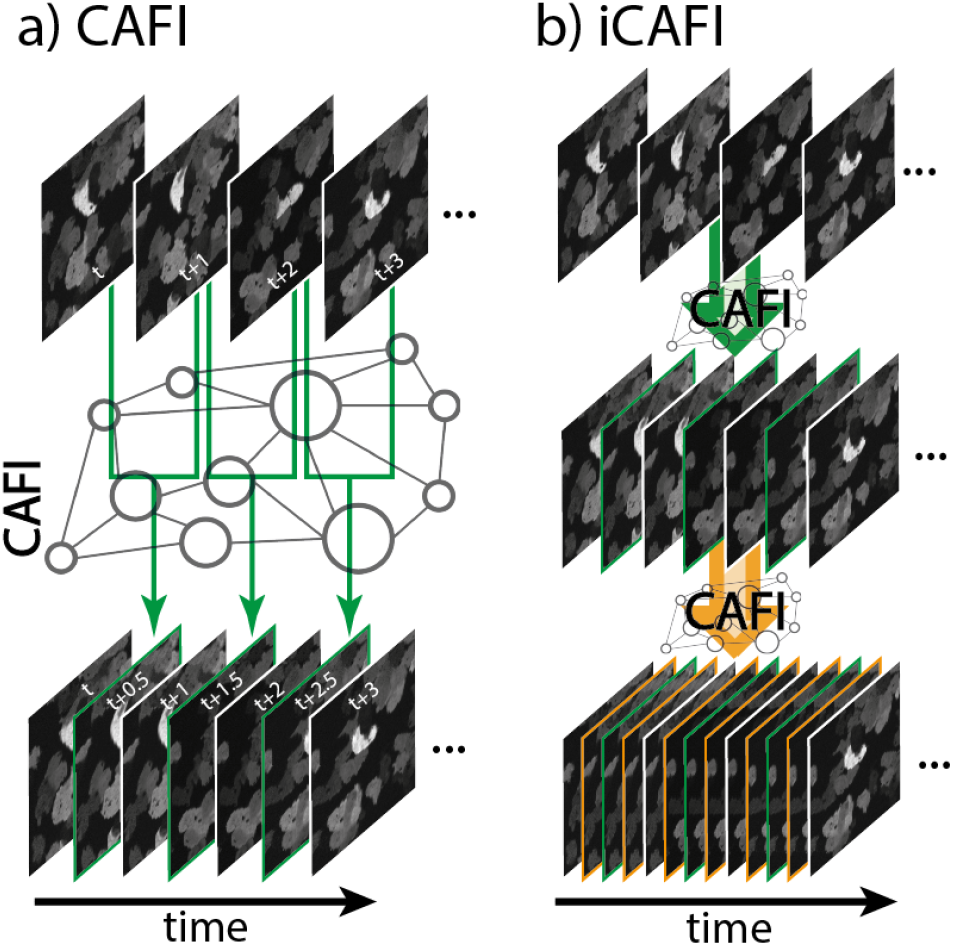
Schematic representation of CAFI. a) CAFI can interpolate images between consecutive frames in a time-course dataset, therefore doubling the resulting temporal frequency. b) iCAFI (iterative CAFI) allows for further improvement of the temporal resolution by repeatedly applying the network prediction.

Deep learning (DL) algorithms have been used for microscopy image post-processing for several years and have transformed the analysis and interpretation of imaging data (2–4). This has led to several breakthroughs for applications in the field of cellular imaging allowing researchers to carry out previously unachievable experiments. For example, DL strategies have been successfully used to improve low SNR images (2, 5, 6), and enhance microscopy image resolution in both the lateral and axial dimensions (7–13).

The computer vision sub-field of video frame interpolation (VFI) developed several DL frameworks for increasing the frame rate of videos to produce slow motion movies (14, 15). The content-aware frame interpolation (CAFI) task is not trivial due to the diversity of the optical flow of the moving objects and frame interpolation neural networks are known to sometimes produce inaccurate predictions or artifacts (15, 16) that could mislead researchers to draw faulty conclusions on biological processes. Despite the great potential of such tools, no fully validated and well-performing CAFI implementations are currently available to the scientific community, let alone as easy-to-use software solutions. Here, we present two implementations of state-of-the-art CAFI networks, Zooming SloMo (ZS) (17, 18) and Depth-Aware Video Frame Interpolation (DAIN) (16) for smart interpolation of microscopy video data. Both networks have shown competitive performance in various benchmark studies against other VFI networks without producing significant visual artifacts on videos (15–18). We demonstrate and compare their abilities to increase the temporal image frequency for a range of microscopy scenarios and perform extensive tracking analysis benchmarks on simulated and real-life experimental data, therefore validating the potentials of the approach.

Surprisingly, even the pretrained models of these networks obtained from natural scenes (moving cars, etc.) perform better than conventional BIC and BIL interpolation. Importantly, we show that fine-tuning these models on appropriate microscopy datasets further improves the quality of the CAFI output. Additionally, we quantitatively demonstrate the improvements achieved by CAFI for the task of single-particle tracking (SPT) using both simulated and an experimental live-cell lysosomal dynamics dataset. Tracking of both datasets showed significant performance improvements after CAFI on several quantitative evaluation criteria.

Overall, we show the improved performance from CAFI over the classical BIL and BIC image interpolation methods on six different datasets from three microscopy modalities (point-scanning confocal, spinning-disk confocal and confocal brightfield microscopy), suggesting that content-awareness can embed relevant information that was not present in the raw dataset. We also demonstrate that CAFI can be used iteratively (iCAFI, see Figure 1b) to increase the restored temporal sampling even further, to 16-fold in the simple case of moving particles. Also, we show that CAFI can be used to improve both the temporal and axial sampling of multidimensional datasets (3D+t) (see Supplementary Figure S1).

We provide the two CAFI network implementations, the corresponding data and pretrained models as part of the Zero-CostDL4Mic platform (19), making CAFI easily available to the wider scientific community both for running predictions and for fine-tuning the networks.

## Results

### CAFI outperforms classical methods in temporal interpolation

To investigate the potential for CAFI to accurately predict time frames, we initially tested the two CAFI networks (DAIN and ZS) on a publicly available mitochondrial dynamics dataset (1). The network predictions were compared with classical interpolation techniques such as BIC, BIL interpolation and simple frame duplication (NONE). The quality of the interpolated images was evaluated using three common objective pixel-based quality metrics: Structural Similarity (SSIM), Root-Mean-Square-Error (RMSE) and Peak-Signal-to-Noise Ratio (PSNR) (2, 20). To objectively compare ZS with DAIN, the ZS network architecture, originally implemented with 4x pixel upsampling (17), was modified to only perform the interpolation without any upsampling. We implemented both networks as ZeroCostDL4Mic notebooks (19) and trained the models using the Vimeo90K-septuplet video dataset (21) (for more details on training see Supplementary Table S1). We hypothesize that training the networks on publicly available and large video datasets, even if unrelated to microscopy, teaches them to recognize general movement dynamics in image sequences that may be useful for live-cell dynamics. For the DAIN network, a pretrained model using this dataset is already available (16). Without any further fine-tuning on microscopy images, both networks already performed significantly better than the classical BIL and BIC interpolation techniques (Figure 2b), demonstrating general dynamics can be learnt from natural scenes and applied to microscopy datasets. After fine-tuning of ZS (FT-ZS) and DAIN (FT-DAIN) models on a subset of the mitochondria dynamics data, the performance of both models further improved (see quality metrics for FT-DAIN and FT-ZS in Figure 2c). Based on the quality metrics, FT-ZS best captured movement patterns of the mitochondria branches, followed by FT-DAIN, and both networks could interpolate movement patterns with greater precision than any classical technique (see Figure 2b, Supplementary Figure S2 and Supplementary Video S1). We noticed that BIL performed better than BIC which created slightly stronger smeared interpolated frames than BIL leading to the weaker performance. As expected, simple frame duplication (NONE) was the worst-performing method as it does not blend or add any new information into the interpolated image frame.

**Fig. 2.**
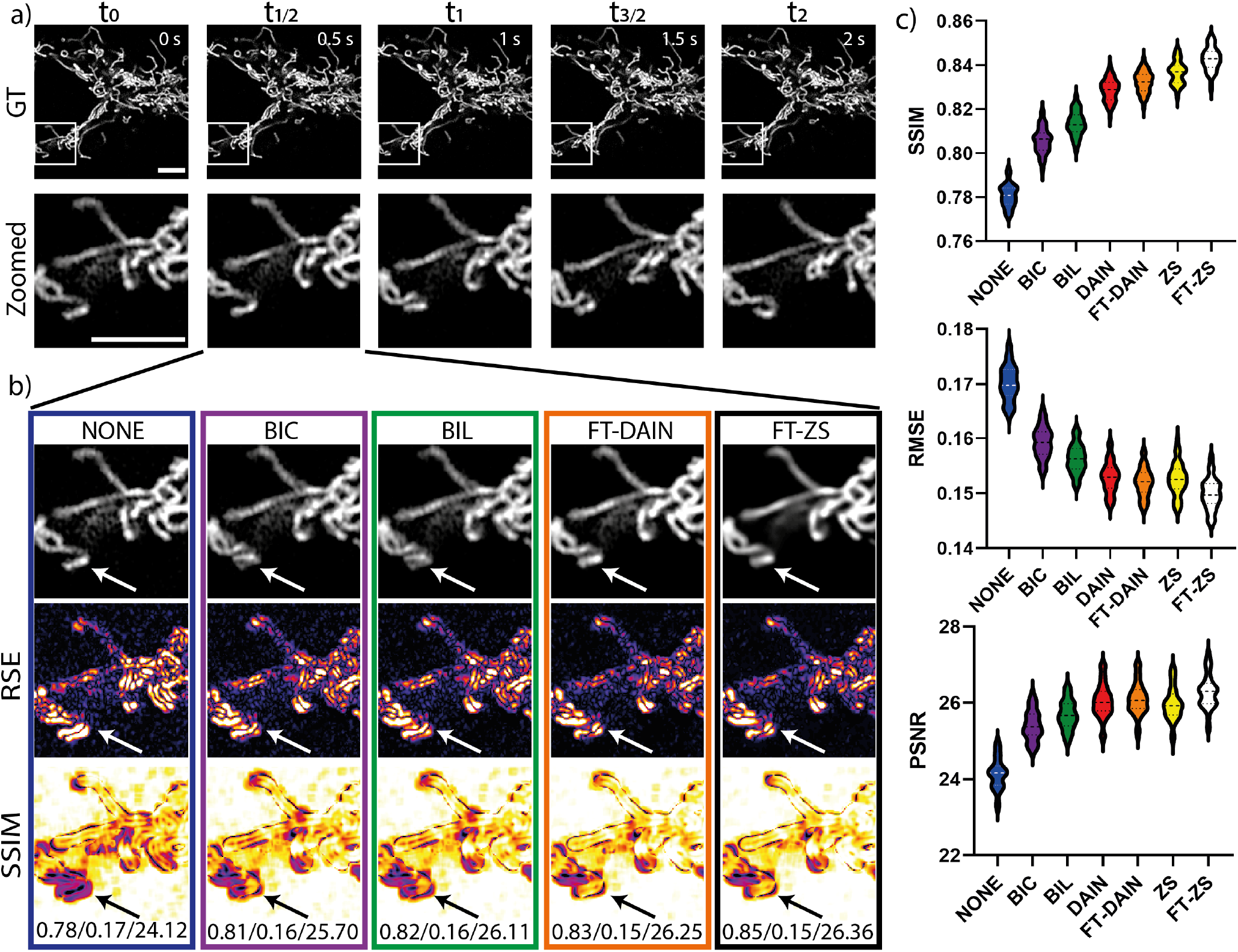
CAFI enables the recovery of fast mitochondrial dynamics. a) Ground truth (GT) image sequence example (top) with zoomed in sections (bottom) of labelled mitochondria branches of U2OS cells (scale bars: 5 µm and timestamp in top right corner of GT images). b) Interpolated frames (top), RSE maps (middle) and SSIM maps (bottom) with white/black arrows highlight areas of fast movement; SSIM/RMSE/PSNR displayed below. c) SSIM, RMSE and PSNR metric results of 49 interpolated image frames compared for each interpolation method including best performing fine-tuned DAIN (FT-DAIN) and Zooming SlowMo (FT-ZS) networks. Data from Fang *et al*. (1).

### Predicting simulated particle motion with CAFI

Lyso-somal behavior is a highly dynamic phenomenon involved in important cellular processes such as degradation and repair mechanisms (22). To study its dynamics, the organelles need to be tracked at speed over long periods of time, which can be a challenge for conventional imaging approaches. To investigate whether CAFI can improve the tracking performance of particle motion such as lysosome dynamics, we initially tested the different interpolation techniques on simulated fluorescent particles moving with a range of motion, as was done for the 2014 ISBI particle tracking challenge (23, 24). The simulated dataset gave us the opportunity to control all parameters of the particle properties such as size, velocity, and contribution of random and directed motion as well as Brownian motion to successfully mimic lysosomal dynamics. Furthermore, the simulated dataset provided the ground truth of the particle locations allowing for easy comparison of the tracking results after CAFI. The full set of parameters used for data generation are given in the Supplementary Table S2 and a detailed explanation on the parameters can be found in the methods section.

First, we evaluated the ability of the networks to cope with increasing particle velocities. For this, ground truth datasets were generated by temporally down-sampling a simulated dataset. An increasing number of frames were removed to simulate increasing particle velocities. The missing frames were subsequently used as ground truth for image quality evaluation. These same missing frames were predicted with BIL interpolation, and DAIN and ZS after fine-tuning the models (see Supplementary Table S1 for further details on training) (see visual illustration of method in Supplementary Figure S3 and for more details on how the data was generated see methods section). The different particle velocities are labeled from V2 to V13 indicating the maximum number of pixels that a particle with directed linear motion could travel from one image to the next (considering that particles not always travel in each frame).

We observed that the interpolation quality of all interpolation techniques decreased with higher particle velocities (see Figure 3a). Also, CAFI clearly outperformed BIL interpolation and ZS tended to perform better than DAIN for low velocities. However, ZS interpolation quality was more significantly affected by increasing velocities than DAIN. This can also be seen qualitatively in Figure 3b and Supplementary Figure S4. Both networks performed very well on particles with slow to medium velocities (see white arrows in Figure 3b (left)) but at higher velocities DAIN performed better than ZS (see white arrows highlighting error regions in Figure 3b (right)). When multiple particles are in close proximity and moving in different directions or when particles move too quickly, DAIN erroneously interpolates the signal in multiple different directions (see error examples in Figure 3c (middle)). ZS did not create any particles if the travelled distance from one frame to the next became too large (see error example in Figure 3c (right)). This observation explains why ZS showed a steeper decrease in image quality at higher movement velocities compared to DAIN. Compared to BIL interpolation both CAFI networks made significantly fewer mistakes (see Figure 3c (left) and statistical analysis in Supplementary Figure S5), again highlighting the power of CAFI for cellular dynamic studies.

**Fig. 3.**
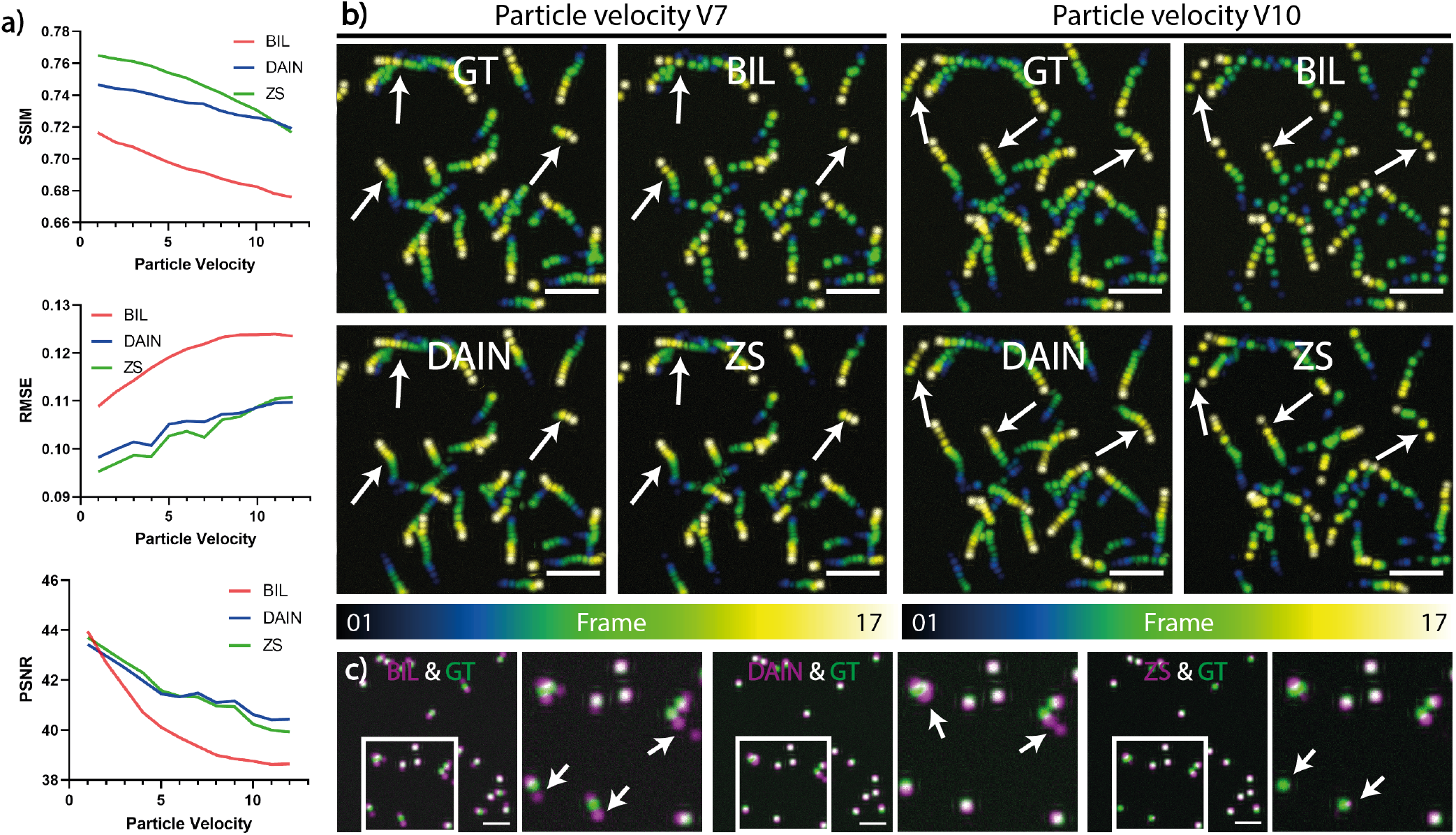
Quantitative assessment of CAFI performance in predicting particle motion using simulated data. a) Image quality metrics comparison (SSIM, RMSE, PSNR) of simulated dataset interpolation with BIL, DAIN and ZS for increasing particle velocities with particle diameters of 15 pixels. b) Temporal color-coded overlaid projection of image sequences of the different interpolation methods for the particle velocities V7 (left) and V10 (right); white arrows highlight regions of interest for comparing interpolation performance (scale bar: 100 pixels). c) Representative artifacts observed from predictions of BIL (left, SSIM: 0.375, RMSE: 0.200, PSNR: 22.58), DAIN (center, SSIM: 0.435, RMSE: 0.187, PSNR: 23.82) and ZS (right, SSIM: 0.455, RMSE: 0.184, PSNR: 23.57) (in magenta) overlaid with ground truth (in green) of the simulated dataset at particle velocity V7. White arrows highlight regions of interest where different techniques make mistakes (scale bars: 50 pixels).

In summary, DAIN and ZS both performed better than BIL for temporal interpolation on the simulated particle datasets. ZS had slightly better performance for slow to moderately moving particles, however ZS made more mistakes for faster object speeds creating blurry artifacts or letting the object disappear. This is where DAIN’s performance is more stable, and it should therefore be used for more challenging and faster moving datasets.

### iCAFI allows for multi-step 16x temporal resolution improvement

A repeated interpolation on the same dataset with a CAFI model has the potential to produce an even higher temporal image frequency from a given dataset (iCAFI, see Figure 1b). However, this approach may lead to an amplification of the errors introduced by the networks in the first interpolation step. To evaluate the performance of iCAFI, we first generated a simulated dataset that was downsampled, removing every second image in 4 iterative steps (2x - 4x - 8x - 16x downsampling). We then explored whether the missing frames can be obtained by iteratively applying BIL, DAIN and ZS predictions (see visual illustration of down- and re-upsampling method in Supplementary Figure S6).

The quality metrics diagrams in Figure 4a show an example for the quality of every frame that was created between two consecutive ground truth image frames (e.g., frame 0 and 16). Based on these quality metrics, both CAFI networks showed very similar performance in the multi-step interpolation. The first interpolation step (predicting the center frame 8) showed the biggest image quality drop, since this was the most demanding interpolation step because of the large travel-distance of the particles from one image to the next. In the following interpolation steps, which consider smaller travel-distances (e.g., frame 0 to 8 or 8 to 16 for the second interpolation iteration) the quality for DAIN and ZS started to recover when approaching the original input images (see in particular U-shape PSNR curve in Figure 4a).

**Fig. 4.**
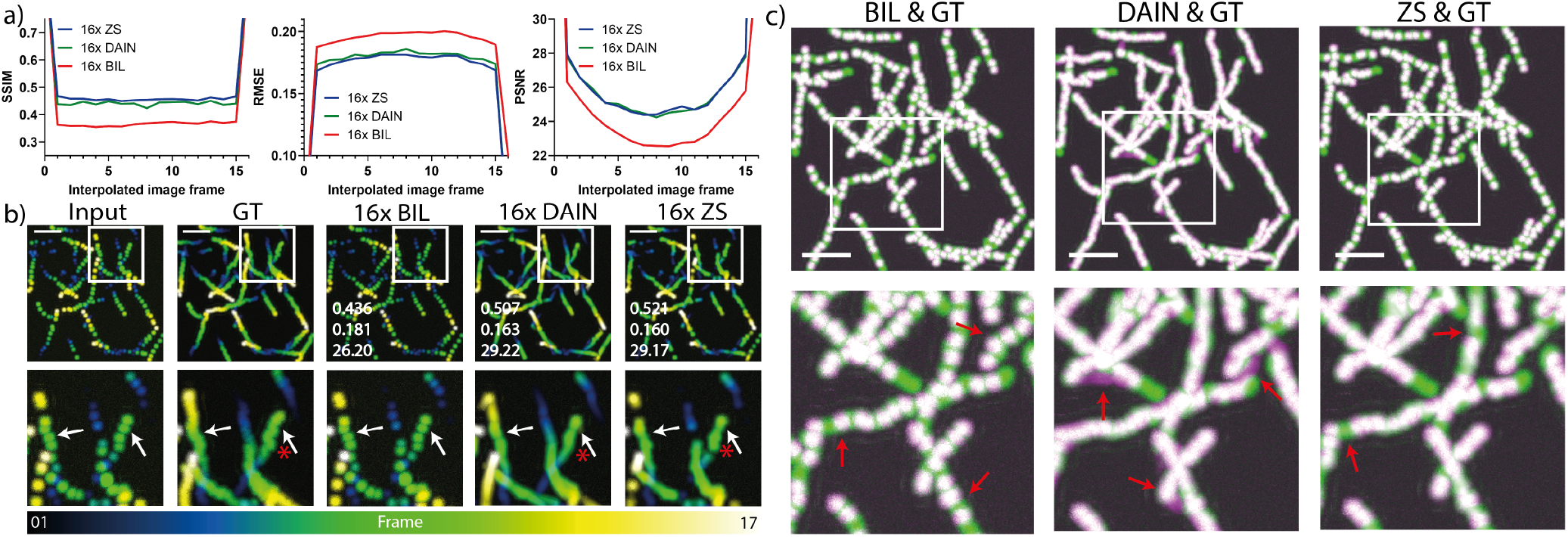
Iterative CAFI (iCAFI) allows for 16-fold accurate interpolation of particle motion prediction. a) Image quality metrics comparison (SSIM, RMSE, PSNR) of BIL, DAIN and ZS on multi-step image interpolation for every image frame between two ground truth input images. b) Temporal color-coded overlaid projection for visual comparison of 16x interpolated image sequences of ground truth (GT), BIL, DAIN and ZS; SSIM/RMSE/PSNR of image sequence shown in overlaid images of each interpolation technique. Red stars highlighting example of Brownian motion loss of CAFI compared to GT. c) Overlaid image-stack projections showing the errors between GT (green) and the interpolated results of BIL, DAIN and ZS (magenta) with matching overlaid parts (white); white boxes indicating zoomed in section and red arrows highlighting regions of interest for error comparison (scale bars: 50 pixels).

By comparing the overlaid maximum intensity temporal image stack projections of DAIN and ZS with the ground truth, the simulated Brownian motion of the particles got lost in the interpolated image sequence for both networks (see red stars in Figure 4b, and demonstration in Supplementary Video S2). However, the direction of the particles can be accurately captured with CAFI (see white arrows in Figure 4b) which was not possible with the BIL interpolation. ZS sometimes missed particles in the first interpolation step that could not be recovered in following interpolation steps (see red arrows in Figure 4c (right)), creating gaps in the predicted trajectory. DAIN, however, captured most particles well but created small artifacts that got amplified in the interpolation steps thereafter (see red arrows Figure 4c (center)). This error amplification explains the slightly worse performance of DAIN compared to ZS for this dataset. Due to the lack of content-awareness, the BIL interpolation was not able to fill the missing spaces between the ground truth input frames of the overlaid temporal projections and therefore performed considerably worse than the two CAFI techniques (see red arrows Figure 4c (left)).

### Quantitative analysis of CAFI performance for single– particle tracking

Pixel-based metrics serve as a useful tool for the computer vision research field in evaluating the performance of model predictions. But before deploying any model for use in scientific research, it is essential to confirm that the downstream analyses (e.g., segmentation, particle tracking statistics) which use the model predictions are sufficiently accurate. To evaluate the quality improvements of the interpolated image sequences obtained from the CAFI networks in the context of SPT, the different particle velocity datasets were used to perform tracking experiments using the TrackMate (25) plugin developed in the image analysis platform Fiji (26). For tracking quality evaluation, the five tracking performance criteria from the ISBI particle tracking challenge were used (23). The different particle velocity tracks obtained from the predictions of the different techniques (BIL, DAIN and ZS) were compared with the actual ground truth position of the tracks (for an explanation of how these tracks were generated see methods section).

At very low particle velocities, all interpolation methods achieved similar results and ZS performed slightly better than DAIN. The tracking results obtained from CAFI predictions outperformed the tracking results of the BIL interpolation by a large margin, especially for intermediate velocities (see V4 to V8 in Figure 5a). The comparison of the tracks confirmed the superiority of CAFI output over those obtained from BIL interpolation. Representative tracking results at velocity V7 are shown in Figure 5b and a representative particle tracking video is provided in Supplementary Video S3.

**Fig. 5.**
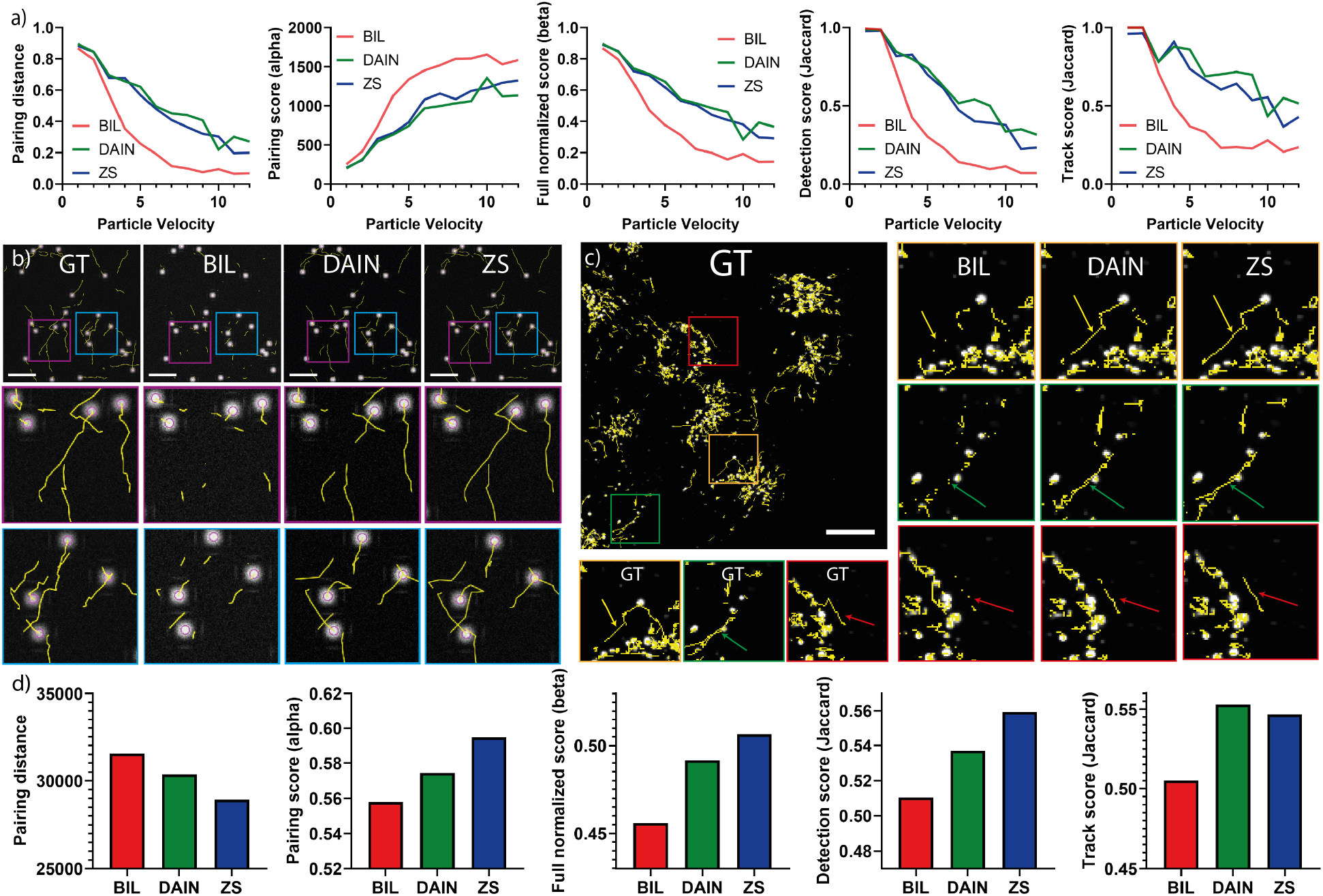
Quautitative assessment of tracking performance of CAFI networks on simulated and experimental data. a) Five tracking evaluation metric results of simulated datasets at different particle velocities with BIL, DAIN and ZS interpolation. b) TrackMate tracks comparison of BIL, DAIN and ZS interpolation compared to ground truth tracks at particle velocity V7 (scale bars: 50 pixels). c) Comparison of lysosomal tracking performance on experimental lysosomal dynamics datasets with zoomed in sections; arrows highlighting region of interest of tracking differences (scale bar: 20 µm). d) Lysosomal tracking performance metrics comparison.

To prove that CAFI can be applied efficiently to real experimental data, we then used CAFI to analyze lysosomal dynamics of live SH-SY5Y cells on a 4D (3D+t) dataset. All z-slices were projected on one image with maximum intensity projection generating a 2D+t dataset. We then removed every other frame to generate a dataset for which ground truth was available. The missing frames from the downsampled dataset were predicted with BIL and CAFI methods and the stacks were then analyzed with TrackMate. The results were compared with the TrackMate tracks of the original (not down-sampled) dataset. On this experimental dataset, both models performed significantly better in all tracking evaluation metrics than the classical BIL interpolation technique. The five evaluation metrics (see Figure 5d) and the comparison of the tracks show clear improvements of CAFI over the classical interpolated tracking results (see Figure 5c and Supplementary Video S4).

### Demonstration of CAFI’s performance on a range of microscopy modalities

After investigating the capabilities of DAIN and ZS to improve tracking results of simulated and experimental datasets, we tested their performance on four more experimental datasets obtained from different microscopy modalities (point-scanning confocal, spinning-disk confocal and confocal brightfield microscopy) to demonstrate the versatility of the CAFI approach. The different datasets tested comprise different motion types such as lysosomal movement, cell migration and fibronectin (27) visualization of different cell lines.

The image sequences were downsampled where every second frame was removed and kept for ground truth comparison and quality evaluation. For each imaging modality we compared the image quality results of CAFI (DAIN and ZS) with and without fine-tuning, BIL, BIC and NONE (frame duplication). The two CAFI networks were each fine-tuned on images of the same microscopy modality (for more details on the training see supplementary Table S1 and methods section). The results of the quality evaluation metrics of all datasets are presented in Table S3. DAIN and ZS trained only on the VIMEO video dataset already outperformed the classical interpolation techniques for all tested datasets. ZS performed better than DAIN in all datasets and after finetuning on the training images of the same imaging modality the prediction quality of both networks further improved. We noted that DAIN sometimes created artifacts by blending the two ground truth image frames together (see arrow in DAIN of Figure 6a and Figure 6d). ZS however, sometimes created washed out and smoothened results where the interpolation details could not be reconstructed with great confidence (see arrow in ZS of Figure 6c). A more detailed interpolation comparisons are shown in Supplementary Figures S7-S10 and video demonstrations of 2x and examples of 4x (iCAFI) frame interpolation are presented in the Supplementary Videos S5-S8.

**Fig. 6.**
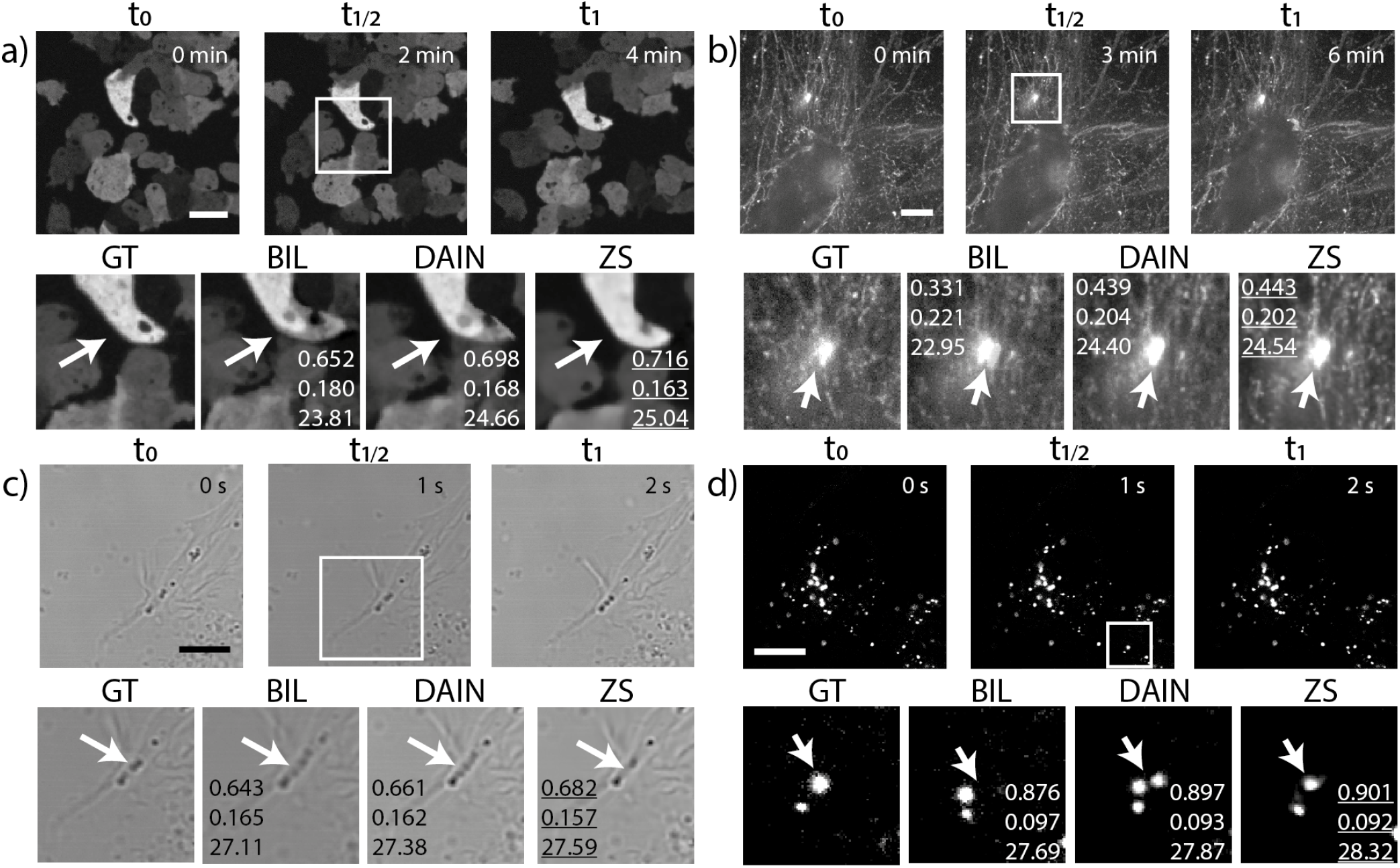
CAFI allows smart interpolation for a wide range of microscopy applications. CAFI interpolation results shown for a) dictyostelium and b) fibronectin labelled A2780 cells (data from Kaukonen *et al*. (27), both recorded on spinning-disk confocal microscopes; c) SH-SY5Y cells recorded with a confocal brightfield microscope; d) labelled lysosomes of SH-SY5Y cells recorded with a confocal microscope; (all scale bars: 10 µm; timestamp in top right corner of GT images); with quality metrics (SSIM/RMSE/PSNR) displayed in zoomed in images. Arrows highlight regions of interest of visible differences between the interpolation techniques.

Given CAFI is ultimately a smart interpolation tool, we also demonstrate that CAFI can perform smart interpolation of the z-stacks on 3D datasets (see Supplementary Figures S11-S12, Supplementary Videos S9-S11 and Supplementary Table S4 for quality comparisons). Additionally, a particularity of ZS is that it is also capable of performing lateral upsampling (see Supplementary Figure S13). We tested this capability and demonstrated its great performance for this task too. For more details see Supplementary Note 2 and demonstrated examples are shown in Supplementary Figures S14-S19 with training conditions and quality comparisons presented in Supplementary Tables S5-S7.

## Discussion

In this work, we present two implementations of state-of-the-art CAFI neural networks (Zooming SlowMo (17, 18) and DAIN (16)) that can effectively increase the frame rate of time-lapse microscopy data by predicting intermediate frames between two consecutive images. Both neural networks showed great performance on mitochondria dynamics (Figure 2) and particle tracking tasks from both simulated and experimental datasets (Figures 3, 4 and 5). We compared their performance to classical interpolation (BIC and BIL) and showed that content-awareness offered better quality images based on a range of image metrics. Furthermore, iCAFI allowed for iterative interpolation of the same dataset allowing for 16x interpolation. The first interpolation step of iCAFI was the most demanding and could lead to artifacts that might be amplified in the following iterations. As expected, we noticed that random movement patterns such as Brownian motion could not be reconstructed using CAFI and these tools should therefore not be used if this is the subject of interest. However, the general directed motion of objects was successfully recovered in a smoothened fashion.

The CAFI networks were also capable of predicting a range of dynamic movement patterns as demonstrated on six different datasets from three different microscopy modalities (point-scanning confocal, spinning-disk confocal, confocal brightfield microscopy). For each dataset both CAFI net-works outperformed classical interpolation techniques such as BIL and BIC interpolation even without fine-tuning of the networks on the images of the same microscopy modality. Their performance increased even further after fine-tuning and both networks were capable of learning and predicting the more complex cellular movement patterns. Furthermore, their strengths and weaknesses in the context of image artifact generation were assessed. For fast moving objects both CAFI networks started to make mistakes and created significant image artifacts, however, for slow and moderate object movement speeds both networks performed very well in their interpolation tasks where DAIN showed more stable results for higher movement speeds and ZS demonstrated greater precision for slower dynamics. ZS created blurry artifacts or missed fast moving objects and DAIN created artifacts by creating movement in different directions.

Finally, DAIN and ZS also outperformed BIL and BIC in the axial interpolation and the lateral upsampling functionalities of ZS was found to achieve good performance too.

These universal content-aware image interpolation solutions show great potential for any microscopy modalities that would benefit from a reduced laser exposure on the sample or a higher frame rate to investigate fast cellular processes. Despite their potential, CAFI networks may still produce image artifacts when imaging fast objects. So, importantly, quality control of model output on appropriate validation data should be performed in the context of downstream analyses (e.g., segmentation, particle tracking) before implementing these tools for real-world investigations, as has been suggested (28). In the future, the simultaneous analysis of multiple channels could provide additional inputs for identifying the dynamics. We also expect that recurrent neural network architectures that have memories for more than the 2 adjacent frames considered here to improve the performance of CAFI approaches, but at the cost of higher computational costs and complexity. The increasing availability of public dataset, especially of dynamic datasets, will also improve the capabilities to build more general models for microscopy and high performance pre-trained models for efficient fine-tuning.

Content-awareness therefore constitutes a powerful approach for microscopy as it facilitates the recovery of high-quality datasets using knowledge embedding acquired at the training stage. As was previously demonstrated for spatial resolution improvement (2), here we show that temporal resolution can also be recovered accurately using CAFI networks such as DAIN and ZS. We expect that this content-awareness may exploit additional context from large multi-dimensional imaging datasets. However, it is important to remember that using content-awareness for image restoration heavily relies on the assumptions that the training dataset fully encompasses all the types of dynamics that will be observed in the analyzed dataset. Deviation from this will almost inevitably lead to errors. So although CAFI provides a powerful tool for live-cell microscopy, it should only be considered when the acquisition of such datasets is not physically possible, and only when the outputs have been validated on downstream workflows using real-world ground truth data.

In summary, here we demonstrate the potentials and versatility of CAFI to improve the frame rate of many microscopy imaging modalities in need for a higher frame frequency. We also provide the tools and pretrained models used in this paper to the wider scientific community through the Zero-CostDL4Mic platform.

## Methods

### Simulated dataset

The simulated dataset was created using the ICY plugin from the 2014 ISBI particle tracking challenge (23, 24). The datasets for temporal interpolation consisted of “switching uniform” white particles moving in front of a black background. The tool allowed to precisely control the image and particle parameters. The image parameters included options for image dimensions, temporal frame density as well as SNR. The parameter related to particle appearance and behavior included particle size, velocity and variables for selecting the contributions of Brownian (sigma) and directed motion fraction. The sigma parameter describes the particle displacement distance of Brownian motion and the directed motion fraction parameters describes the probability for Brownian motion and directed motion to occur. The parameter of “xy pixel size” was chosen to generate particles with diameters of roughly 15 pixels which was a good size estimate for real-life lysosomes recorded with 63x magnification. The movement related parameters were empirically selected to simulate the behavior of lysosomal dynamics most accurately. All parameter values for the particle simulation datasets are provided in Supplementary Table S2.

For the different velocity experiments one densely framed image sequence (225 frames) was generated. To generate the different ground truth velocities an increasing number of images for each velocity dataset was removed between each two consecutive time point frames (1 to 12 removed images were labelled as velocities V2 to V13). Therefore, each dataset had a gradually increased particle travel distance from one frame to the next and therefore also an increased overall particle velocity. These ground truth velocity datasets where then downsampled by removing every second image before re-interpolating them with the provided different interpolation techniques (see visual illustration of this preparation methodology in Supplementary Figure S3). Furthermore, the ground truth image sequences were limited to 17 frames to make them easily comparable.

For the iCAFI multi-step temporal interpolation experiment, the same densely framed simulated dataset (225 frames) was downsampled, removing every second image in four iterative steps (2x - 4x - 8x - 16x downsampling), while keeping the removed images as ground truth for comparison for the following iterative re-interpolation steps (see Supplementary Figure S6 for visual illustration of the down- and re-upsampling process). As training data for fine-tuning of the CAFI networks 10 simulated datasets with 225 images per sequence were created with the same parameters as the test data, including the full equivalent range of velocities to be predicted (equivalent to velocities V2 to V13).

### Particle tracking

Particle tracking was performed using the Fiji plugin TrackMate (25). For detecting the simulated dataset particles, the LoG detector with an estimated blob diameter of 10 and a threshold of 2 was used. The linking step was performed with the “Simple LAP tracker” with a linking distance of 15 - 20 µm, gap-closing of 15 - 20 µm depending on the particle velocity and maximum frame gap of 2. For the full set of parameters for the different particle tracking scenarios as well as for the real-life lysosomes tracking experiment can be found in Supplementary Table S8 and S9.

### Particle speed ground truth tracks generation

The ground truth tracks for the different particle velocities were generated in the following steps. First the big, simulated dataset (225 frames) was generated with the ICY data generator. This plugin provided a XML file with all the ground truth time points and precise particle point locations. This XML file was first converted into the ISBI XML format in the TrackMate interface and the relevant time points of this file was downsampled in the same way as the actual image sequences, where an increasing number of frames was removed between each time point (see visual illustration of method in Supplementary Figure S3). A developed python script selected just the time points and particle coordinates relevant for the specific movement velocity and generated a new XML file containing just the locations of the particles for the selected frames in that particle velocity option. These ground truth tracks were then compared with the TrackMate detected tracks of the BIL and the CAFI interpolated image sequences and the five performance criteria from the ISBI particle tracking challenge (23) were evaluated with the associated ICY ISBI Challenge Tracking Batch Scoring plugin for comparing the different tracking files.

The ground truth tracks for the real-life lysosomal data were evaluated as the TrackMate generated tracks of the full image sequence before downsampling of the dataset. The evaluated tracks from TrackMate were exported in the ISBI challenge format. Then the TrackMate tracks of the downsampled and re-interpolated image sequences were compared to their ground truth tracks using the ISBI Challenge Tracking Batch Scoring plugin from ICY (23).

### Network training

The ZS interpolation with the options of 1x (no lateral upsampling) and 2x lateral upsampling was initially trained from scratch on the Vimeo90K-septuplet (21) (82GB) dataset. First the full dataset was split into smaller 8-12 GB sub-datasets which was necessary because of quota limitations of the Google Colab environment. Then the training was carried out for a total of 300 epochs for DAIN and 600.000 niter for ZS. Before using the two CAFI models for the different interpolation tasks both networks were fine-tuned with 0.5 to 14 GB of images of the same imaging modality which was depended on the amount of available training data. The learning rate for training of both networks was reduced to 1e^-5^ while the other parameters were used as provided from the original papers (16, 17). For more details on the training datasets and additional parameters see Supplementary Table S1.

For the training of the lateral upsampling functionality, ZS and SRFBN-S were fine-tuned on 3.6 to 12.1 GB training data with a learning rate of 1e^-5^ and 1e^-4^, respectively. For fine-tuning of ZS for 4x lateral upsampling of the electron microscopy dataset to compare it with the PSSR network, gaussian noise was added to the provided training data as it was done in the original paper (1) (for more details on the datasets and the epoch/niter sizes see Supplementary Table S5).

To increase the amount of available training data of the Dictyostelium dataset the recorded images with a resolution of 1200×1200 pixels were augmented by zooming to each corner part of the image with the image size of 512×512 pixels and by resizing to the full target image down to 512×512 pixels.

The data preparation for small datasets was performed in the provided Google Colab notebooks. For datasets bigger than 2 GB the training data preparation was performed offline with the python data preparation scripts provided in the github repository. The created folders where then uploaded on Google Drive/Google Cloud Storage for training in the Google Colab environment.

### Lysosome cell imaging

SH-SY5Y cells were cultured in Dulbecco’s Modified Eagle Medium (DMEM, Invitrogen, Carlsbad, CA) supplemented with 10% Fetal Bovine Serum (FBS, Invitrogen), glutamine (2 mM), and penicillin/streptomycin (50 µg/mL, Invitrogen). All cells were grown in a 5% CO2 incubator at 37 °C. The cells were plated and grown on 8-well chamber slides (LabTek II Chamber Coverglass) in 250 µL of culture media at a plating density of 25,000 cells per well and allowed to grow for 24 h. Next the media was changed to media containing lipofectamine 2000 (2 µL/mL) and the lysosomal copper probe FLCS1 (60 nM) (29). The cells were incubated with this dye for 24 h. Prior to imaging the cell media was changed back to DMEM with 10% FBS (250 µL/well). The image acquisition was performed on a confocal microscope (Leica SP5) with 63x magnification 1.4 NA oil objective. 3D image (XYZT) time series were recorded for 20 minutes collecting 40 time points of 30 z-stack images at a resolution of 512px (0.481 µm/px) and temporal time sequences (XYT) were recorded with the same settings at a imaging speed of 1 seconds/frame.

### Dictyostelium Imaging

Dictyostelium discoideum cells were genetically engineered to transiently express eGFP and td-tomato from an extrachromosomal plasmid and grown adherently in HL5 medium. Prior to imaging, cells were washed in KK2 medium and transferred onto KK2-agar pads at a density of approximately 5 × 10^5^ cells cm^-2^. Once cells had adhered to the pads, the pads were inverted onto ibidi u-dishes and overlaid with silicone oil to prevent dehydration. Samples were imaged on an inverted spinning-disk confocal microscope (3i), using a 63x 1.4NA oil objective and equipped with a prime95B CMOS camera (photometrics). 3D image (XYZT) time series were recorded with 2-minute frame intervals.

### DAIN architecture

The Depth-Aware Video Frame Interpolation (DAIN) was published by Wenbo *et al*. in 2019 (16). This network was trained to detect occlusions by exploring depth information and based on this information it performs the frame interpolation. The network learns hierarchical features by gathering contextual information from neighboring pixels. The interpolated frame is synthesized by combining the information in an adaptive wrapping layer by integrating the two input frames, depth maps and contextual features based on optical flow and a local interpolation kernel. The depth maps functionality from the input frames is performed by an hourglass network (a special type of convolutional encoder-decoder network) pretrained on the MegaDepth dataset (30). Furthermore, the flow estimation is performed by a pretrained PWC-Net (31) and the contextual information is obtained by using a pretrained ResNet architecture (32). A U-Net network is then used for the kernel estimation and an adaptive wrapping layer combines all the information flows from each sub-network. To ensure that the network predicts residuals between the ground truth frame and the blended frame, the two warped frames are linearly blended. The pretrained network used for transfer learning was trained on the Vimeo90K dataset (21) (82GB) which is a large-scale, high-quality video dataset consisting of 89,800 video clips downloaded from the VIMEO streaming platform. For a more detailed explanation about the architecture of the network see the original paper (16).

### Zooming SlowMo architecture

The Zooming SlowMo network is based on a paper published by Xiaoyu Xiang *et al*. in 2021 (17). The network can perform next to the image frame interpolation (VFI) also a simultaneous video super-resolution (VSR) increasing the image resolution up to 4-times in the same processing step. The Zooming SlowMo network allows for a one-stage process of directly reconstructing high-resolution and high frame rate image sequences. The network uses a deformable feature interpolation network to get feature-level temporal information and combines it with a deformable ConvLSTM to aggregate the temporal information. This allows for handling of motions and effectively leveraging global contexts with simultaneous temporal alignment and aggregation. The Zooming SlowMo network consists of four main parts which are a feature extractor, frame feature temporal interpolation module, deformable ConvLSTM, and an HR frame reconstructor. The feature extractor with a convolution layer first produces feature maps which are then used to synthesize LR intermediate frames in the frame feature interpolation module. Then the ConvLSTM performs a simultaneous alignment and aggregation for the consecutive feature maps. In the last step the HR sequence is constructed from the aggregated feature maps. For a more detailed explanation about the architecture of the network see the original paper (17).

## Code availability

The CAFI source code and documentation (of Zooming SlowMo and DAIN) are available for download on Github, https://github.com/mpriessner/CAFI and are free for non-profit use.

## Data availability

Example training data and pretrained models are included in the GitHub release (v1.0.0). Our training and testing data sets are made available via Zenodo https://zenodo.org/record/5596603.YX-bKGDMIdU.

## ACKNOWLEDGEMENTS

M.P. is thankful to the UK Engineering and Physical Sciences Research Council (EPSRC) for a studentship as part of the Centre of Doctoral Training Neurotechnology (EP/L016737/1). U.M., and A.S. are supported by the Waitt Foundation and NIH-NCI P30 Grant No. 014195 and by NSF NeuroNex Award No. 2014862. U.M. is supported by the Chan-Zuckerberg Initiative Imaging Scientist Award. R.F.L. would like to acknowledge the support of the MRC Skills development fellowship (MR/T027924/1). The Facility for Imaging by Light Microscopy (FILM) at Imperial College London is part-supported by funding from the Wellcome Trust (grant 104931/Z/14/Z) and BBSRC (grant BB/L015129/1). T.L. and J.R.C. are supported by the Wellcome Trust: Wellcome Trust Award (UK) 202867/Z/16/Z

