## Supplementary information for "Content-aware frame interpolation (CAFI): Deep Learning-based temporal super-resolution for fast bioimaging"

### Supplementary Note 1: Links to ZeroCostDL4Mic Platform and to Google Colab notebooks

CAFI Github page:
https://github.com/mpriessner/CAFI
ZeroCostDL4Mic Wiki:
https://github.com/HenriquesLab/ZeroCostDL4Mic/wiki
DAIN 4 Microscopy Notebook:
https://colab.research.google.com/drive/1bL6wgTWrghHK7LH9xb4KGSk5WuOa5nJS?usp=sharing
ZoomingSlowMo 4 Microscopy Notebook:
https://colab.research.google.com/drive/1TZ0K-rq9Nrgu9_XZ0UOK6brxjIM0ISNU?usp=sharing

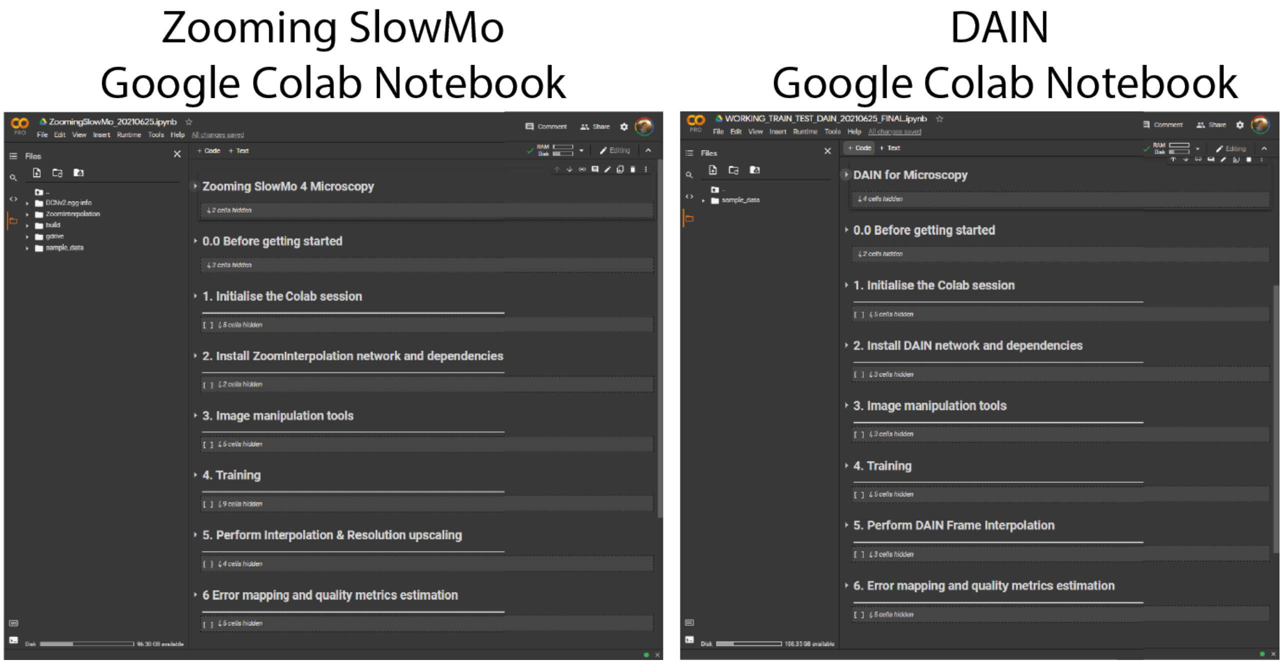


Screenshots of the two ZeroCostDL4Mic Google Colab notebooks for DAIN and ZS.

### Supplementary Note 2: Image upscaling using Zooming SlowMo CAFI network

On top of ZS' interpolation functionality, it also has the ability to simultaneously increase the lateral image resolution while performing the temporal or axial content-aware interpolation. The 2x and 4x lateral upsampling capabilities of ZS were compared with BIL and BIC upsampling and one other lateral upsampling neural network called SRFBN-S which showed state-of-the-art competitive results in the so called "Image super-resolution" upsampling task (33). The ZS and SRFBN-S neural networks were fine-tuned on training data of the same imaging modality (for more details on the training see Supplementary Table S5). ZS clearly outperforms BIL and BIC upsampling and creates better results compared to the SRFBN-S neural network (see Supplementary Figures S14-S19). Finally, we compared the lateral upsampling functionality of ZS with the 4x upsampling results of the provided noisy electron microscopy images of the recently published PSSR network (1). Here ZS was fine-tuned on the training data with added gaussian noise as it was done in the original paper for PSSR. ZS produced clearly better results compared to BIC/BIL and also equivalent results compared to PSSR (see examples in Supplementary Figure S20 and delta comparisons in Supplementary Figure S21). The full quality evaluation results (SSIM, RMSE, PSNR) for 2x and 4x lateral upsampling are presented in Supplementary Table S6 and S7.

### Supplementary Videos

**Supplementary Video S1:** Demonstration of CAFI networks for temporal interpolation on confocal microscopy dataset of fluorescently labelled mitochondria branches (data from Fang *et al.* (1)).

**Supplementary Video S2:** Demonstration of CAFI networks for temporal interpolation of a simulated particles dataset generated with the ISBI particle tracking challenge plugin from Icy (24). Particle diameter is 15 pixels and the selected movement velocity is ranging from V2 to V8.

**Supplementary Video S3:** Demonstration of TrackMate (25) tracking improvements of simulated particles generated with the ISBI particle tracking challenge plugin from Icy (24) after temporal interpolation with CAFI. Particle diameter is 15 pixels and the movement velocity selected ranging from V2 to V8.

**Supplementary Video S4:** Demonstration example of lysosomal tracking improvements after temporal interpolation with CAFI using TrackMate (25). Lysosomes were labelled with FLCS1 (29) and the image sequence was collected as 3D+t dataset on a Leica SP5 with a 63x magnification 1.4 NA oil objective. Z-stacks were projected with maximum intensity generating a 2D+t dataset. The tracking results from the full image sequence was taken as ground truth tracks for quality comparison.

**Supplementary Video S5:** Demonstration of CAFI networks for temporal interpolation on inverted spinning-disk confocal microscopy dataset of GFP labelled dictyostelium cells. Images were recorded using a 63x 1.4NA oil objective with 2-minute frame intervals.

**Supplementary Video S6:** Demonstration of CAFI networks for temporal interpolation on a spinning-disk confocal microscopy dataset of fluorescently labelled fibronectin of A2780 cells (data from Kaukonen *et al.* (27)).

**Supplementary Video S7:** Demonstration of CAFI networks for temporal interpolation on confocal brightfield dataset of SH-SY5Y cells. Images were recorded on a Leica SP5 with a 63x magnification 1.4 NA oil objective with 1-second frame intervals. The image sequence was down-sampled removing every second image in two iterative steps and re-interpolated in temporal dimension with 2x CAFI and 4x iCAFI.

**Supplementary Video S8:** Demonstration of CAFI networks for temporal interpolation on a point-scanning confocal microscopy dataset of fluorescently labelled lysosomes of SH-SY5Y cells. Images were recorded on a Leica SP5 with a 63x magnification 1.4 NA oil objective with 1-second frame intervals. The image sequence was down-sampled removing every second image in two iterative steps and re-interpolated in temporal dimension with 2x CAFI and 4x iCAFI.

**Supplementary Video S9:** Demonstration of zCAFI networks for axial interpolation on an electron microscopy dataset of rat hippocampus (data from Fang *et al.* (1)). The image sequence was down-sampled removing every second image in two iterative steps and re-interpolated in axial dimension with 2x CAFI and 4x iCAFI.

**Supplementary Video S10:** Demonstration of zCAFI networks for axial interpolation on a structured illumination microscopy (SIM) dataset of fluorescently labelled actin of DCIS.COM cells (data from Weigert *et al.* (2)).

**Supplementary Video S11:** Demonstration of tzCAFI on fluorescently labelled fibronectin on a 4D spinning-disk confocal microscopy dataset (data from Kaukonen *et al.* (27)). The networks were trained for the interpolation task on data in the temporal dimension and the same fine-tuned network was used for both interpolation dimensions (axial and temporal).

### Supplementary Tables and Figures

| Training / fine tuning for image frame interpolation | | | | | | | |
|---|---|---|---|---|---|---|---|
| **Dataset** | **Microscopy type** | **Test dim** | **Pixel dimension** | **Size** | **epochs DAIN\|niter ZS** | **LR DAIN\|ZS** | **Batch size DAIN\|ZS** |
| **VIMEO video dataset** | - | T | 488x256 | 82 GB | 300\|600000 | 1e-5\|1e-5 | 1\|16 |
| **Mitochondria Data** | **Confocal microscope** | T | 512x512 | 14 GB | 10\|20000 | 1e-5\|1e-5 | 1\|16 |
| **Dictyostelium Data** | **Spinning disc confocal microscope** | T | 512x512 | 2.6 GB | 40\|60000 | 1e-5\|1e-5 | 1\|16 |
| **SH-SY5Y Data** | **Brightfield microscope** | T | 512x512 | 3.6 GB | 30\|50000 | 1e-5\|1e-5 | 1\|16 |
| **Synthetic Data** | - | T | 512x512 | 450 MB | 40\|80000 | 1e-5\|1e-5 | 1\|16 |
| **Lysosomal Data** | **Confocal microscope** | T | 512x512 | 5.2 GB | 20\|40000 | 1e-5\|1e-5 | 1\|16 |
| **Fibronectin Data** | **Spinning disc confocal microscope** | T/Z | 512x512 | 300 MB | 40\|80000 | 1e-5\|1e-5 | 1\|16 |
| **Actin Data** | **Structured illumination microscope** | Z | 512x512 | 2.4 GB | 20\|30000 | 1e-5\|1e-5 | 1\|16 |
| **Hippocampus Data** | **Electron microscope** | Z | 512x512 | 1 GB | 50\|80000 | 1e-5\|1e-5 | 1\|16 |

**Table S1.** Training data and parameter information for fine-tuning DAIN and ZS networks for the image interpolation task of the different datasets. The other network parameters were kept as default.

| ICY (ISBI Challenge Track Generator) | |
|---|---|
| | **T-Dimension Dataset** |
| **SNR:** | 10 |
| **Density:** | 40 |
| **Image width:** | 512 |
| **Image height:** | 512 |
| **Image depth:** | 1 |
| **Sequence length:** | 225 |
| **MinTrackLength:** | 52 |
| **Warmup length:** | 50 |
| **Extinction rate:** | 0.003 |
| **x borders simu:** | 50 |
| **y borders simu:** | 50 |
| **z borders simu:** | 20 |
| **xy pixel size:** | 15 |
| **Slice spacing:** | 250 |
| **Creator types:** | SWITCHING_UNIFORM |
| **Sigma Brownian:** | 1 |
| **q1:** | 2 |
| **V min:** | 2 |
| **V max:** | 2 |
| **Probe directed Brownian:** | 0.3 |
| **Probe Brownian directed:** | 0.3 |

**Table S2.** Parameter table for creating simulated datasets with ICY plugin (34).

| 2x temporal interpolation | | | | | | | | | | | | | | | | | | | | | | | |
|---|---|---|---|---|---|---|---|---|---|---|---|---|---|---|---|---|---|---|---|---|---|---|---|
| | | | 2x-NONE | | | 2x-BIC | | | 2x-BIL | | | 2x-DAIN | | | 2x-DAIN (FT) | | | 2x-ZS | | | 2x-ZS (FT) | | |
| | Dim | Image nr. | SSIM | RMSE | PSNR | SSIM | RMSE | PSNR | SSIM | RMSE | PSNR | SSIM | RMSE | PSNR | SSIM | RMSE | PSNR | SSIM | RMSE | PSNR | SSIM | RMSE | PSNR |
| **Dictyostelium Data** | T | 51 | 0.744 | 0.160 | 26.750 | 0.816 | 0.121 | 37.107 | 0.815 | 0.124 | 35.815 | 0.856 | 0.096 | 42.981 | 0.856 | 0.096 | 43.007 | 0.866 | 0.095 | 42.517 | **0.867** | 0.094 | 42.917 |
| **SH-SY5Y Data** | T | 63 | 0.778 | 0.089 | 78.786 | 0.800 | 0.107 | 40.953 | 0.803 | 0.114 | 37.792 | 0.828 | 0.101 | 42.051 | 0.829 | 0.101 | 42.057 | 0.832 | 0.100 | 41.429 | **0.835** | 0.099 | 42.113 |
| **Synthetic Data** | T | 113 | 0.748 | 0.104 | 41.014 | 0.762 | 0.112 | 35.975 | 0.766 | 0.114 | 36.499 | 0.808 | 0.095 | 42.598 | 0.810 | 0.095 | 42.619 | 0.818 | 0.094 | 42.651 | **0.820** | 0.093 | 42.555 |
| **Lysosomal Data** | T | 57 | 0.922 | 0.049 | 80.319 | 0.926 | 0.061 | 41.127 | 0.924 | 0.060 | 40.900 | 0.935 | 0.046 | 81.321 | 0.940 | 0.045 | 81.237 | 0.941 | 0.047 | 53.423 | **0.942** | 0.046 | 54.979 |
| **Mitochondrial Data** | T | 97 | 0.891 | 0.101 | 40.834 | 0.899 | 0.107 | 35.580 | 0.902 | 0.107 | 35.827 | 0.914 | 0.094 | 41.203 | 0.916 | 0.094 | 41.230 | 0.918 | 0.094 | 41.150 | **0.921** | 0.093 | 41.310 |
| **Fibronectin Data** | T | 13 | 0.663 | 0.108 | 78.749 | 0.680 | 0.128 | 42.939 | 0.682 | 0.130 | 42.188 | 0.732 | 0.104 | 46.883 | 0.733 | 0.104 | 46.896 | 0.732 | 0.103 | 46.617 | **0.734** | 0.104 | 45.067 |

**Table S3.** Quality evaluation of NONE, BIC, BIL, DAIN, FT-DAIN, ZS and FT-ZS interpolation in temporal dimension on 6 different datasets. The numbers in red and blue indicate the best and second-best performance.

| 2x axial interpolation | | | | | | | | | | | | | | | | | | | | | | | |
|---|---|---|---|---|---|---|---|---|---|---|---|---|---|---|---|---|---|---|---|---|---|---|---|
| | | | 2x-NONE | | | 2x-BIC | | | 2x-BIL | | | 2x-DAIN | | | 2x-DAIN (FT) | | | 2x-ZS | | | 2x-ZS (FT) | | |
| | Dim | Images nr. | SSIM | RMSE | PSNR | SSIM | RMSE | PSNR | SSIM | RMSE | PSNR | SSIM | RMSE | PSNR | SSIM | RMSE | PSNR | SSIM | RMSE | PSNR | SSIM | RMSE | PSNR |
| **Fibronectin Data** | Z | 13 | 0.679 | 0.123 | 40.273 | 0.714 | 0.128 | 36.798 | 0.715 | 0.128 | 36.422 | 0.732 | 0.112 | 41.386 | 0.733 | 0.111 | 41.409 | 0.701 | 0.114 | 41.025 | **0.752** | 0.108 | 41.653 |
| **Actin Data** | Z | 33 | 0.769 | 0.127 | 35.265 | 0.813 | 0.126 | 33.219 | 0.807 | 0.130 | 32.395 | 0.810 | 0.105 | 40.077 | 0.812 | 0.105 | 40.105 | 0.812 | 0.103 | 40.180 | **0.870** | 0.093 | 41.984 |
| **Hippocampus Data** | Z | 97 | 0.748 | 0.134 | 39.966 | 0.775 | 0.146 | 34.249 | 0.780 | 0.148 | 33.565 | 0.801 | 0.121 | 41.449 | 0.802 | 0.120 | 41.483 | 0.806 | 0.119 | 41.587 | **0.811** | 0.118 | 41.698 |

**Table S4.** Quality evaluation of NONE, BIC, BIL, DAIN, FT-DAIN, ZS and FT-ZS interpolation on 3 different datasets in axial dimension. The numbers in red and blue indicate the best and second-best performance.

| Training / fine tuning for lateral upscaling | | | | | | | | |
|---|---|---|---|---|---|---|---|---|
| **Dataset** | **Microscopy type** | **Upscale factor** | **LR pixel dimension** | **HR pixel dimension** | **Size HR** | **epochs SRFBN\|niter ZS** | **LR SRFBN\|ZS** | **Batch size SRFBN\|ZS** |
| **VIMEO video dataset** | - | **2x** | 488x256 | 244x128 | 82 GB | - \|600000 | - \|1e-5 | - \|16 |
| **VIMEO video dataset** | - | **4x** | 488x256 | 122x64 | 82 GB | - \| provided | - \| provided | - \| - |
| **SH-SY5Y Data** | **Confocal brightfield microscope** | **2x** | 1024x1024 | 512x512 | 3.6 GB | 10\|20000 | 1e-4\|1e-5 | 16\|16 |
| **SH-SY5Y Data** | **Confocal brightfield microscope** | **4x** | 1024x1024 | 512x512 | 3.6 GB | 10\|20000 | 1e-4\|1e-5 | 16\|16 |
| **Dictyostelium Data** | **Spinning disc confocal microscope** | **2x** | 512x512 | 256x256 | 4.3 GB | 10\|20000 | 1e-4\|1e-5 | 16\|16 |
| **Dictyostelium Data** | **Spinning disc confocal microscope** | **4x** | 512x512 | 128x128 | 4.3 GB | 10\|20000 | 1e-4\|1e-5 | 16\|16 |
| **Hippocampus Data** | **Electron microscope** | **2x** | 512x512 | 256x256 | 12.1 GB | 5\|10000 | 1e-4\|1e-5 | 16\|16 |
| **Hippocampus Data** | **Electron microscope** | **4x** | 512x512 | 128x128 | 12.1 GB | 5\|10000 | 1e-4\|1e-5 | 16\|16 |
| **Lysosomal Data** | **Confocal microscope** | **2x** | 512x512 | 256x256 | 5.2 GB | 10\|20000 | 1e-4\|1e-5 | 16\|16 |
| **Lysosomal Data** | **Confocal microscope** | **4x** | 512x512 | 128x128 | 5.2 GB | 10\|20000 | 1e-4\|1e-5 | 16\|16 |

**Table S5.** Training data and selected parameters for fine-tuning ZS and SRFBN-S networks for the lateral upsampling task for the different datasets. The other network parameters were kept as default.

| 2x lateral upsampling | | | | | | | |
|---|---|---|---|---|---|---|---|
| **Dataset** | **Method** | **LR** | **HR** | **slice** | **SSIM** | **RMSE** | **PSNR** |
| **SH-SY5Y Data** | **ZS** | 256 | 512 | 11 | **0.9658** | 0.0912 | 38.493 |
| | **SRFBN-S** | 256 | 512 | 11 | 0.9634 | 0.0925 | 38.135 |
| | **BIC** | 256 | 512 | 11 | 0.9564 | 0.1016 | 35.605 |
| | **BIL** | 256 | 512 | 11 | 0.9249 | 0.1200 | 32.631 |
| **Dictyostel. Data** | **ZS** | 256 | 512 | 13 | **0.7281** | 0.1568 | 29.978 |
| | **SRFBN-S** | 256 | 512 | 13 | 0.7256 | 0.1571 | 29.932 |
| | **BIC** | 256 | 512 | 13 | 0.7054 | 0.1584 | 29.780 |
| | **BIL** | 256 | 512 | 13 | 0.6779 | 0.1609 | 29.481 |
| **Lysosomal Data** | **ZS** | 256 | 512 | 11 | **0.7714** | 0.1432 | 26.270 |
| | **SRFBN-S** | 256 | 512 | 11 | 0.7631 | 0.1429 | 26.259 |
| | **BIC** | 256 | 512 | 11 | 0.7132 | 0.1500 | 25.492 |
| | **BIL** | 256 | 512 | 11 | 0.6889 | 0.1539 | 25.023 |

**Table S6.** Quality evaluation metrics (SSIM, RMSE, PSNR) for 2x lateral upsampling. The numbers in red and blue indicate the best and second-best performance, respectively.

.

| 4x lateral upsampling | | | | | | | |
|---|---|---|---|---|---|---|---|
| **Dataset** | **Method** | **LR** | **HR** | **slice** | **SSIM** | **RMSE** | **PSNR** |
| **SH-SY5Y Data** | **ZS** | 256 | 1024 | 11 | **0.8473** | 0.1212 | 33.943 |
| | **SRFBN-S** | 256 | 1024 | 11 | 0.8423 | 0.1223 | 33.725 |
| | **BIC** | 256 | 1024 | 11 | 0.8252 | 0.1281 | 32.491 |
| | **BIL** | 256 | 1024 | 11 | 0.7855 | 0.1394 | 30.737 |
| **Dictyostel. Data** | **ZS** | 128 | 512 | 13 | **0.6193** | 0.1676 | 28.736 |
| | **SRFBN-S** | 128 | 512 | 13 | 0.6188 | 0.1678 | 28.720 |
| | **BIC** | 128 | 512 | 13 | 0.6015 | 0.1702 | 28.407 |
| | **BIL** | 128 | 512 | 13 | 0.5762 | 0.1733 | 28.018 |
| **Lysosomal Data** | **ZS** | 128 | 512 | 11 | 0.6367 | 0.1615 | 24.109 |
| | **SRFBN-S** | 128 | 512 | 11 | **0.6375** | 0.1612 | 24.102 |
| | **BIC** | 128 | 512 | 11 | 0.6227 | 0.1658 | 23.455 |
| | **BIL** | 128 | 512 | 11 | 0.6110 | 0.1692 | 23.069 |
| **Hippo. Data** | **ZS** | 125 | 500 | 42 | **0.4085** | 0.2413 | 22.659 |
| | **PSSR** | 125 | 500 | 42 | 0.4076 | 0.2414 | 22.648 |
| | **BIC** | 125 | 500 | 42 | 0.3797 | 0.2603 | 21.371 |
| | **BIL** | 125 | 500 | 42 | 0.3980 | 0.2528 | 21.861 |

**Table S7.** Quality evaluation metrics (SSIM, RMSE, PSNR) for 4x lateral upsampling. The numbers in red and blue indicate the best and second-best performance, respectively. The EM lateral upsampling results were just available for the comparison with PSSR. SRFBN-S failed on this dataset due to mishandling of the introduced noise in the training data.

| TrackMate Parameters Simulated Dataset | |
|---|---|
| **Estimated blob diameter:** | 10 µm |
| **Threshold:** | 2 |
| **Selected Tracker:** | Simple LAP tracker |
| **Linking max distance:** | 15 (V1-6); 20 (V7-10); 25 (V11-12) |
| **Gap-closing max distance:** | 15 (V1-6); 20 (V7-10); 25 (V11-12) |
| **Gap-closing max frame gap:** | 2 |

**Table S8.** Parameter table for tracking of simulated particles with Fiji Trackmate plugin (25).

| TrackMate Parameters Lyso-Dataset | |
|---|---|
| **Estimated blob diameter:** | 3 µm |
| **Threshold:** | 10 |
| **Selected Tracker:** | Simple LAP tracker |
| **Quality particles** | Number calibrated to GT number |
| **Linking max distance:** | 8 |
| **Gap-closing max distance:** | 8 |
| **Gap-closing max frame gap:** | 2 |

**Table S9.** Parameter table for tracking of lysosomal particles with Fiji Trackmate plugin (25).

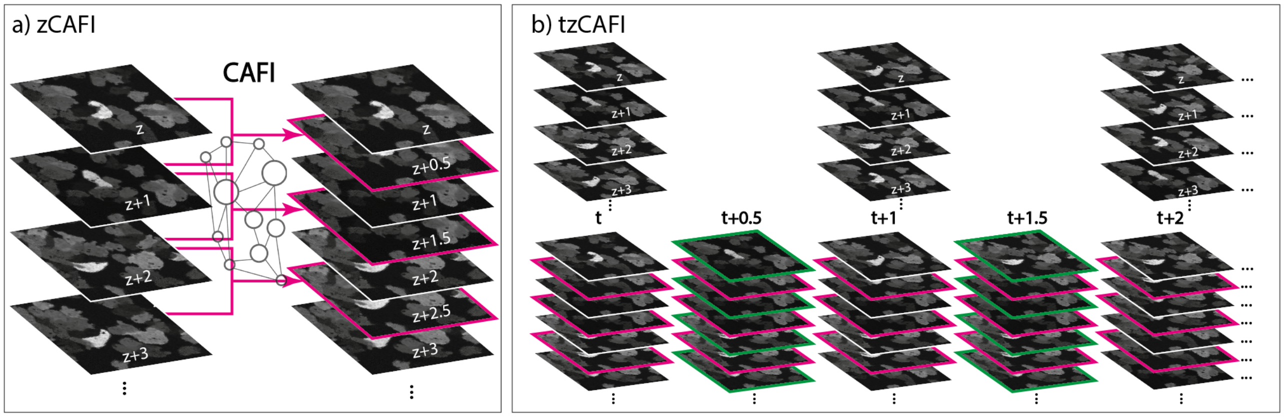


**Fig. S1.** Schematic representation of two more CAFI functionalities. a) zCAFI interpolates images between consecutive images along the z-axis of a dataset. b) tzCAFI sequentially applies both zCAFI (a) and CAFI (Figure 1a main text) on 4D (3D+t) dimensional dataset.

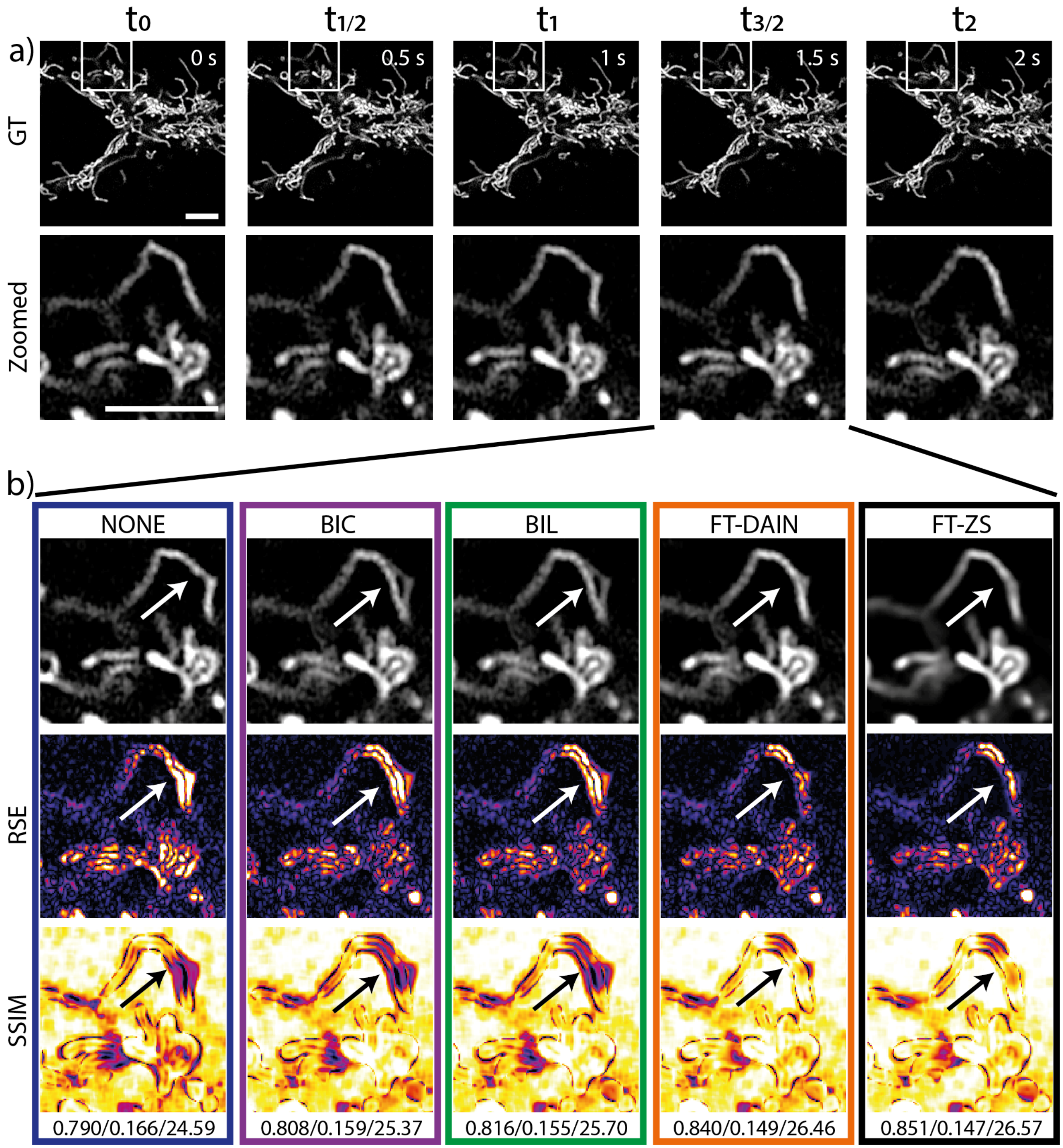


**Fig. S2.** 2x frame interpolation of mitochondrial dynamics image sequence of U2OS cells using classical interpolation with BIL, BIC, NONE, and CAFI with DAIN and ZS. a) Ground truth (GT) image sequence example (top) with zoomed in sections (bottom) (scale bar: 5 µm; timestamp in top right corner of GT images). b) Interpolated frame (top), RMSE map (middle) and SSIM map (bottom) with white/black arrows highlighting movement regions; SSIM/RMSE/PSNR displayed below. Data from Fang *et al.* (1).

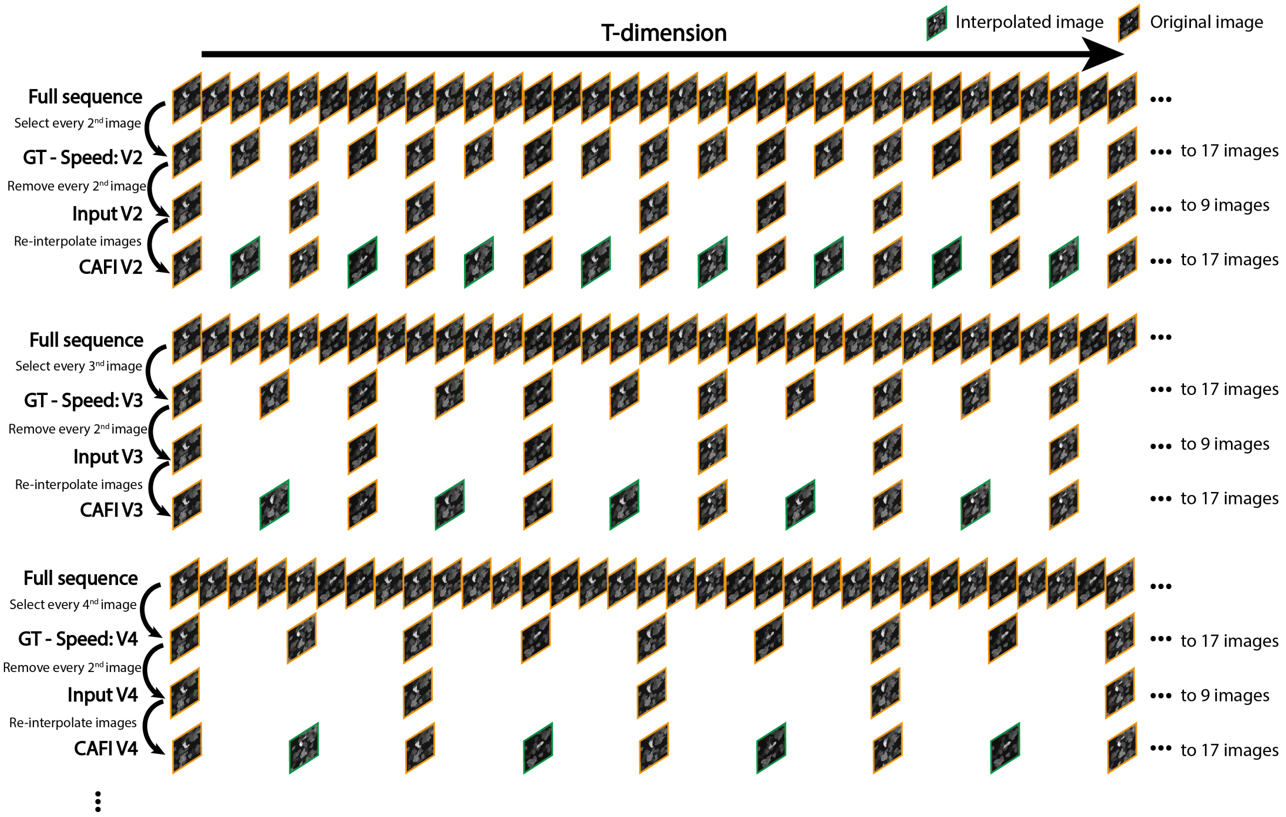


**Fig. S3.** Visual illustration of downsampling process to generate the ground truth (GT) for the different particle velocities. By removing an increasing number of images between two time points increased particle velocities are created as new GT datasets. Then every second image is removed and kept for quality evaluation and is then re-interpolated with the different interpolation techniques.

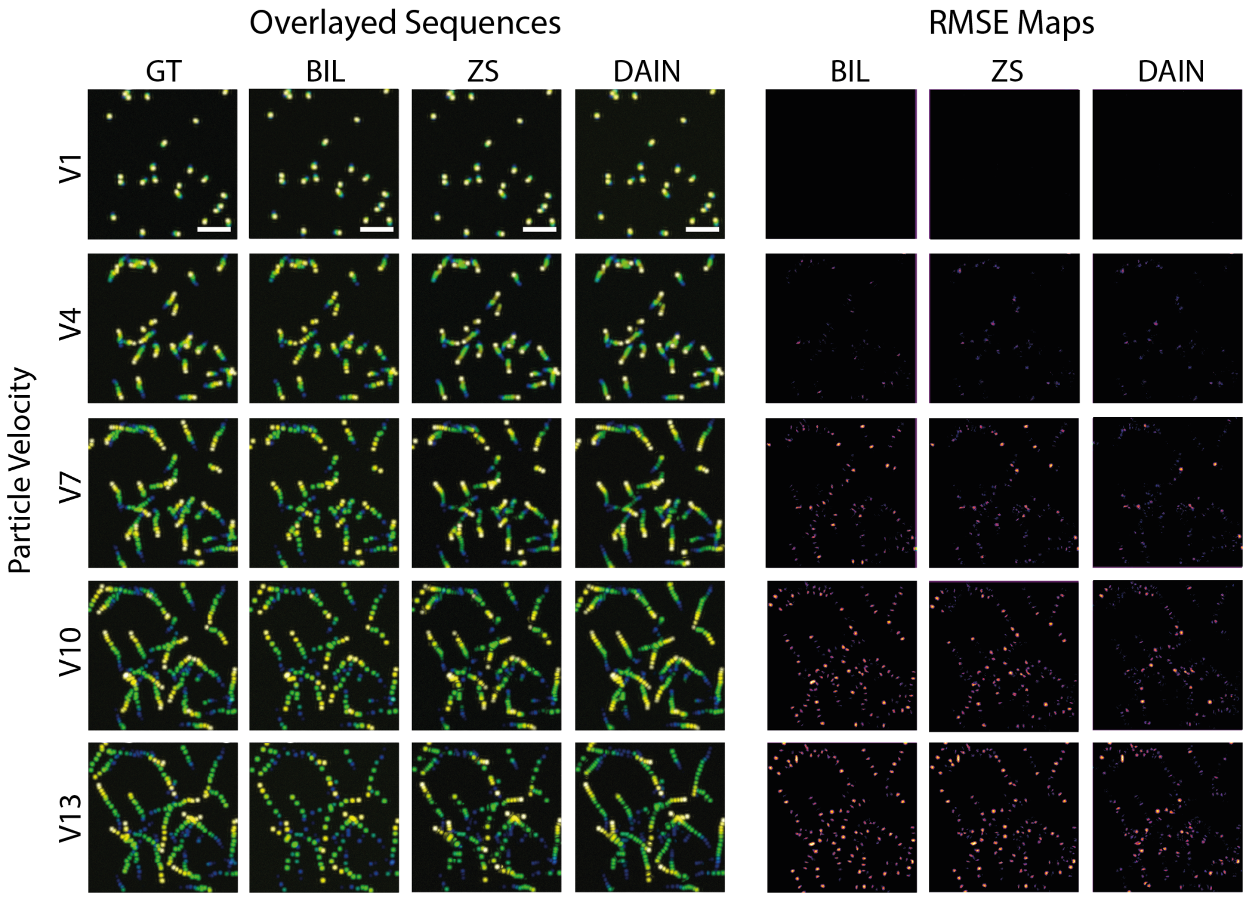


**Fig. S4.** Maximum intensity projection of the simulated time sequence dataset of the different image frame interpolation results (BIL, DAIN, ZS) at different particle movement velocities (left) with RMSE maps compared to ground-truth (right). Scale bars: 50 pixels.

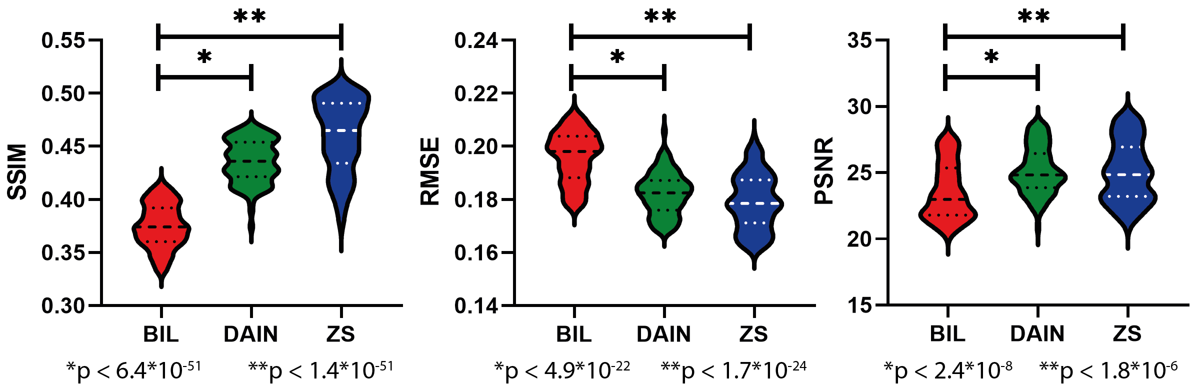


**Fig. S5.** Statistical analysis (two-sided student t-test) of the image quality metrics (SSIM, RMSE, PSNR) by comparing the values of DAIN and ZS of each interpolated image for all particle velocities with the quality metric results of BIL, showing strong significance of improvements for DAIN and ZS over BIL interpolation.

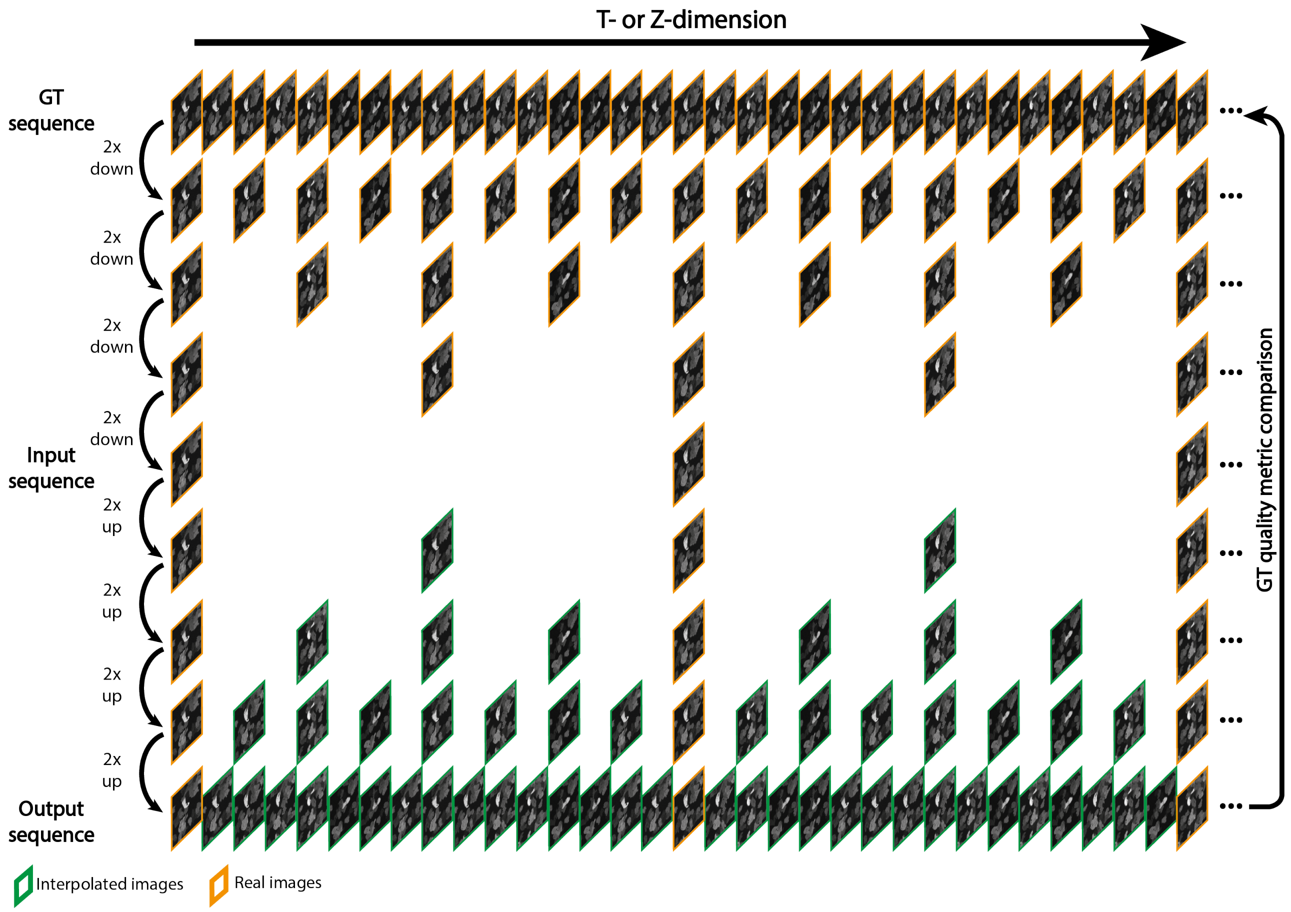


**Fig. S6.** Visual illustration of multi-step frame down- and upsampling for iCAFI. In several iterative steps every second image gets removed and is later re-interpolated in several steps reconstructing the high temporal frequency of the ground truth image sequence.

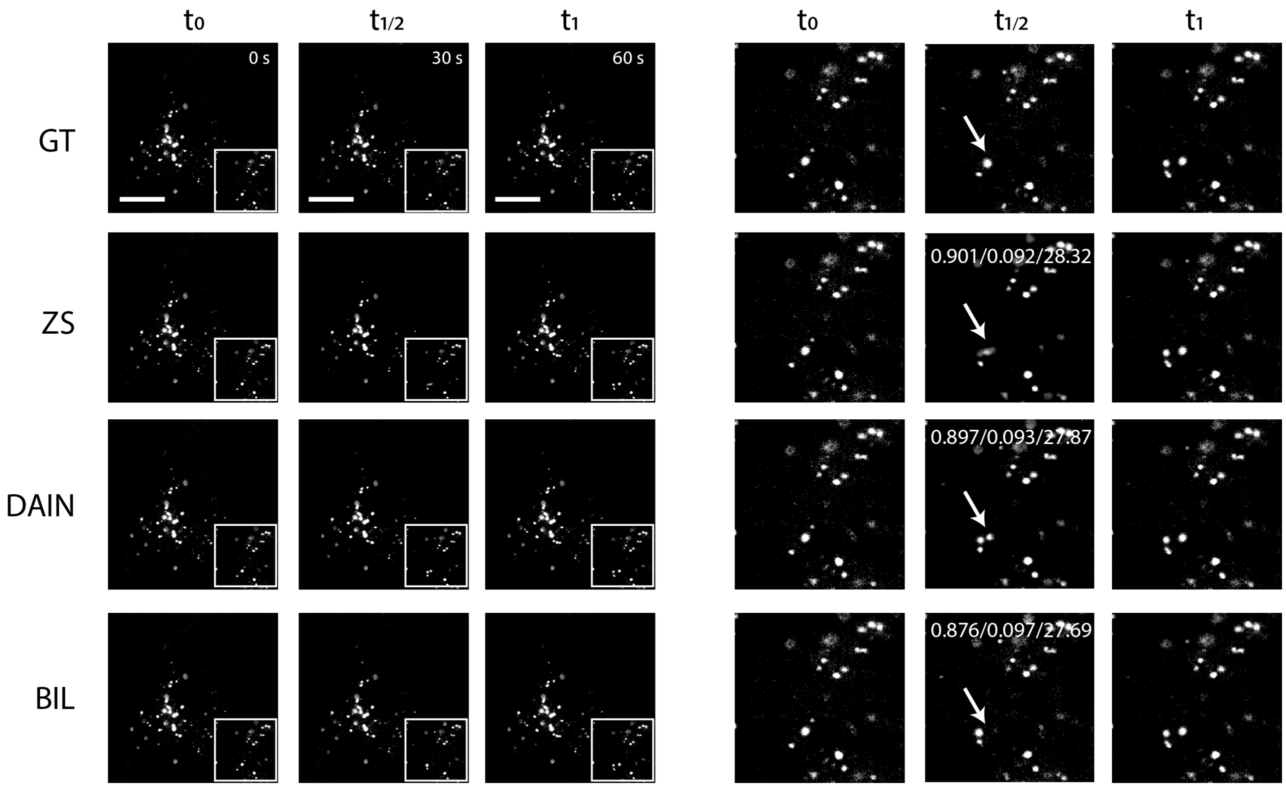


**Fig. S7.** 2x frame interpolation of BIL, DAIN and ZS for lysosomes of SH-SY5Y cells recorded on a confocal microscope. Overall ZS produces slightly better-quality results than DAIN even though ZS misses faster moving lysosomes which DAIN is able to capture well (example highlighted with white arrow in zoomed in sections on the right). SSIM/RMSE/PSNR values displayed in the zoomed in interpolated images. CAFI tools significantly outperform BIL interpolation. Scale bar: 10 µm.

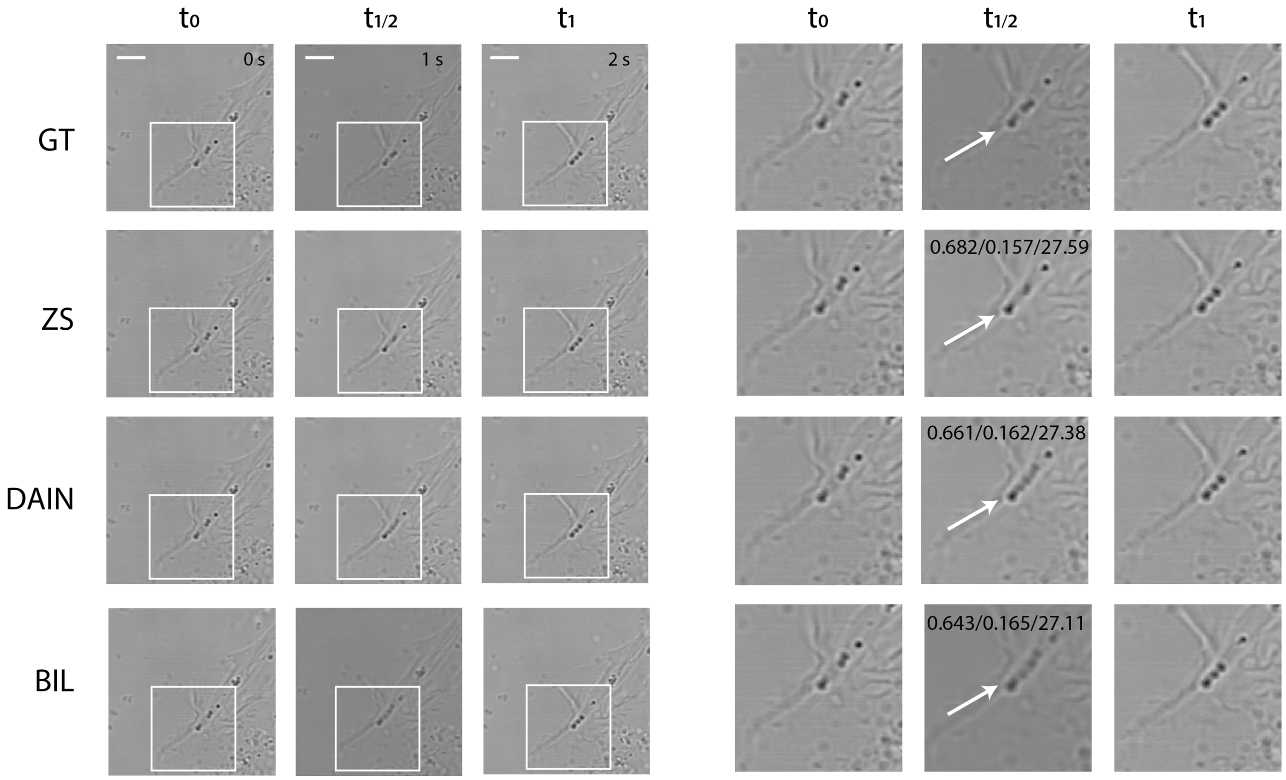


**Fig. S8.** 2x frame interpolation with BIL, DAIN and ZS for SH-SY5Y cells recorded on a confocal brightfield microscope. ZS produces slightly better-quality results than DAIN. DAIN creates visual artifacts for fast moving lipid droplets while ZS manages to capture the movement better (see white arrow in zoomed in sections on the right). SSIM/RMSE/PSNR values displayed in the zoomed in interpolated images. CAFI tools significantly outperform BIL interpolation. Scale bar: 10 µm.

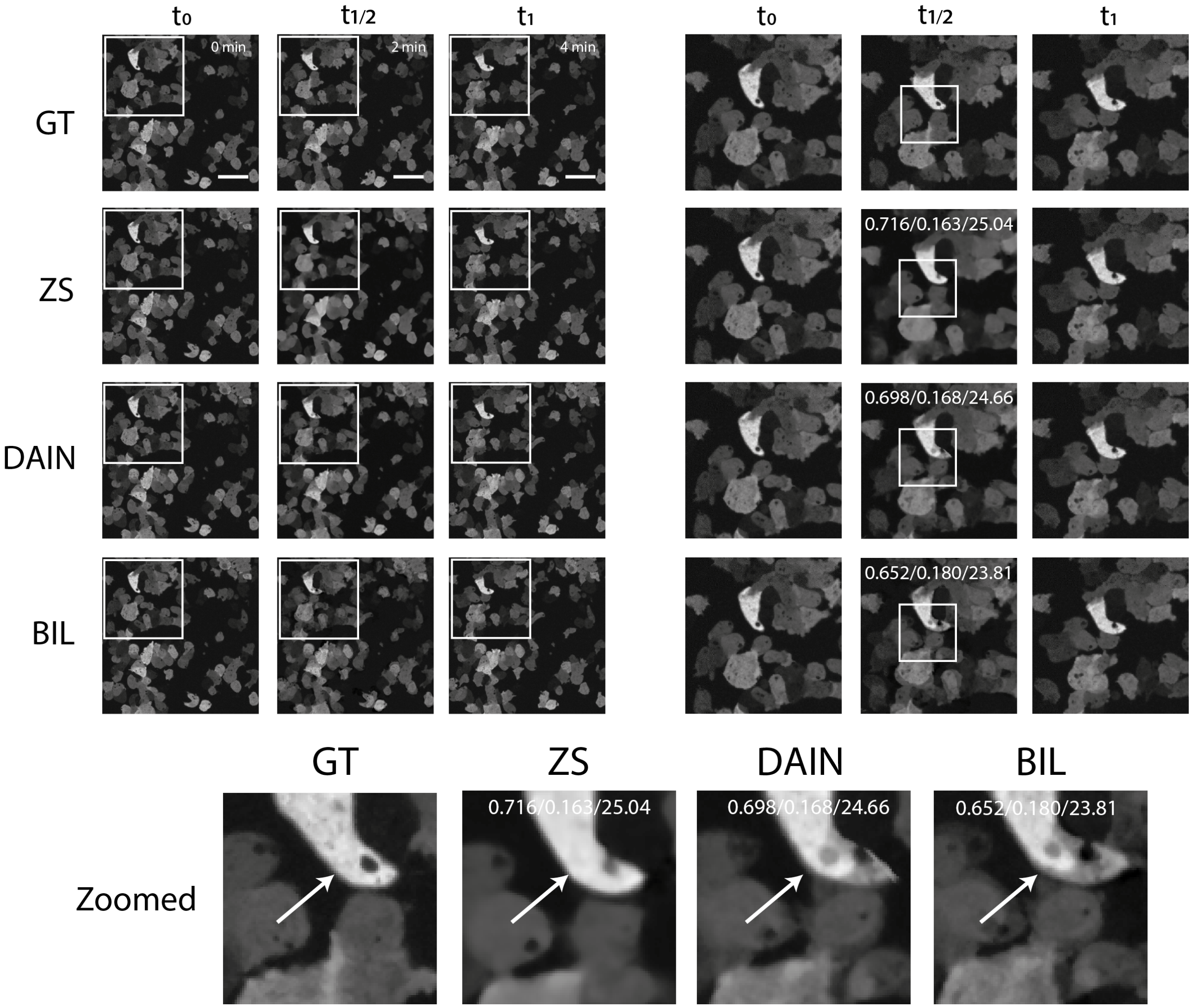


**Fig. S9.** 2x image frame interpolation of BIL, DAIN and ZS for dictyostelium recorded on a spinning-disk confocal microscope. ZS creates smoother transitions and better-quality interpolation results than DAIN (see white arrow in zoomed in section). SSIM/RMSE/PSNR values displayed in the zoomed in interpolated images. CAFI tools significantly outperform BIL interpolation. Scale bar: 20 µm.

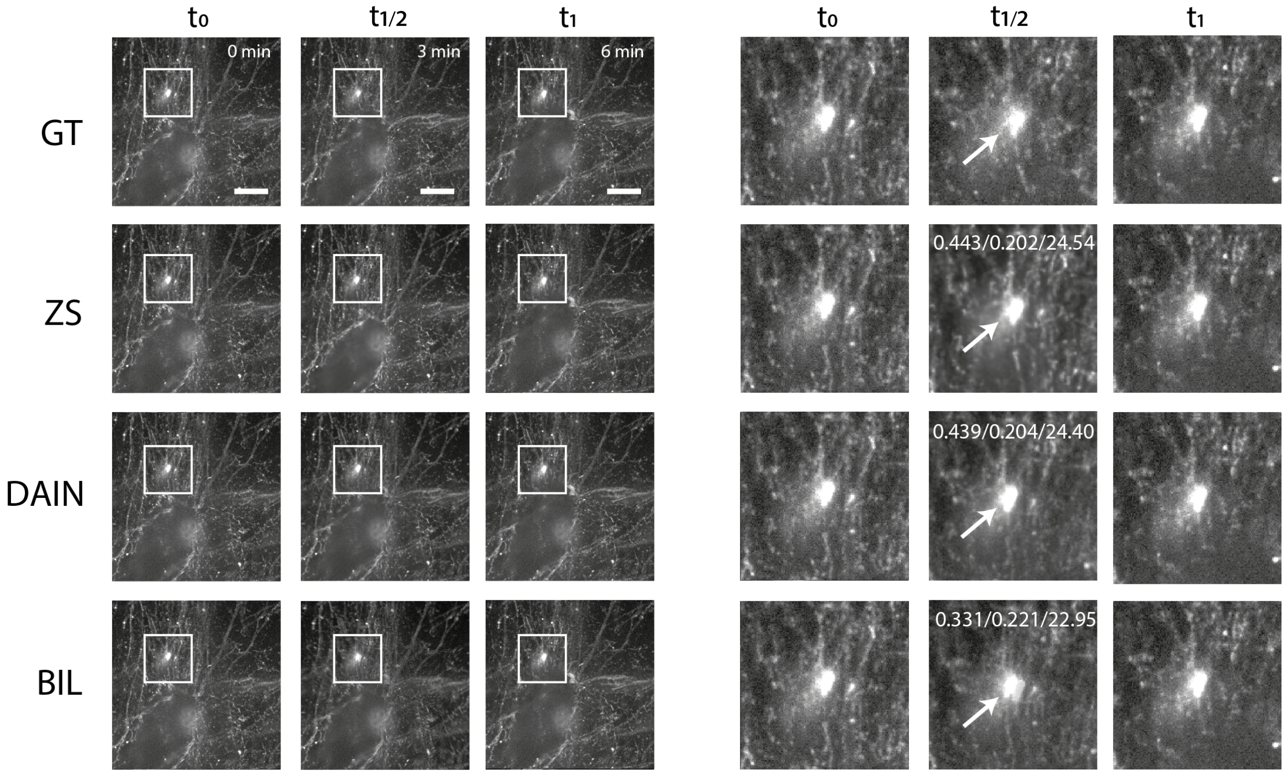


**Fig. S10.** 2x image frame interpolation of BIL, DAIN and ZS for fibronectin labelled A2780 cells recorded on a spinning-disk confocal microscope. ZS creates smoother transitions and better-quality interpolation results than DAIN (see white arrow in zoomed in sections on the right). SSIM/RMSE/PSNR values displayed in the zoomed in interpolated images. CAFI tools significantly outperform BIL interpolation. Scale bar: 10 µm. Data from Kaukonen *et al.* (27).

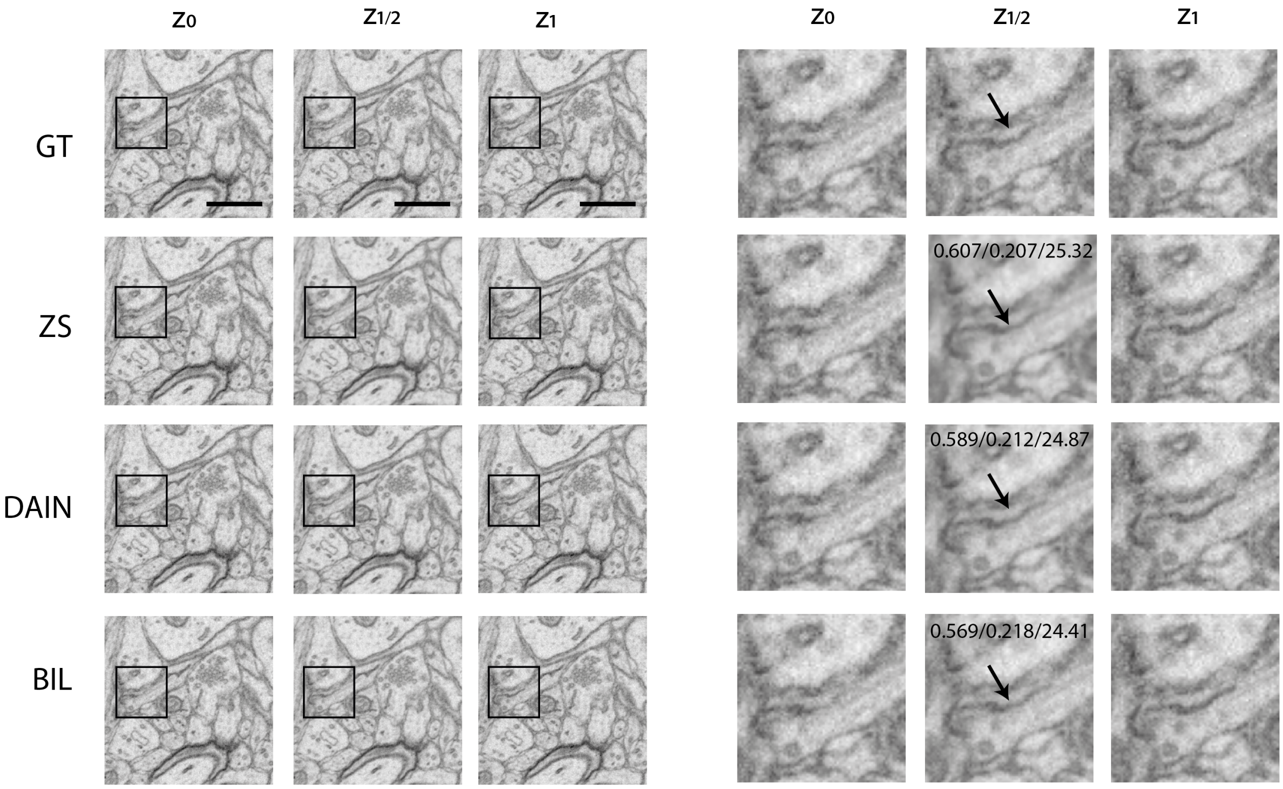


**Fig. S11.** 2x image slice interpolation in axial dimension with BIL, DAIN and ZS for rat hippocampus tissue recorded on an electron microscope. ZS produces better quality results than DAIN. ZS creates smoother transitions of an imaged dendrite (see black arrow in zoomed in sections on the right). SSIM/RMSE/PSNR values displayed in the zoomed in interpolated images. CAFI tools significantly outperform BIL interpolation. Scale bar: 0.4 µm. Data from Fang *et al.* (1).

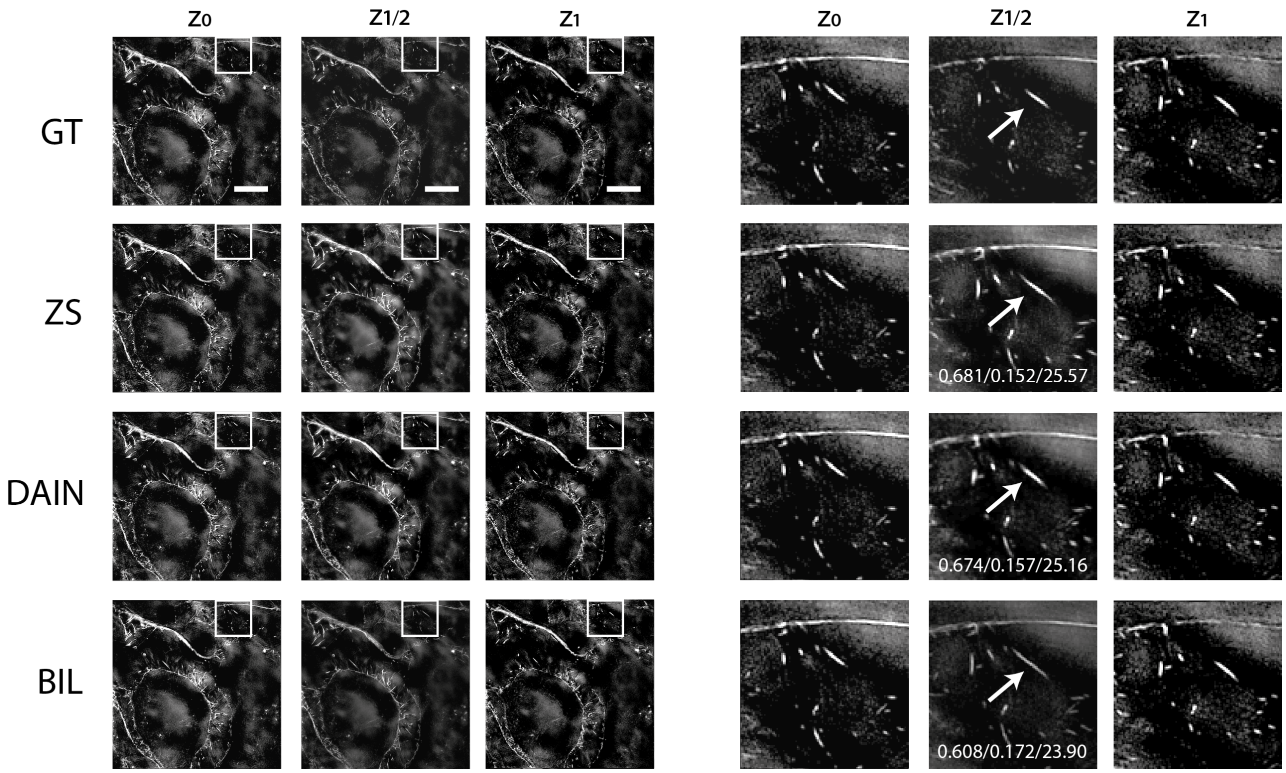


**Fig. S12.** 2x image slice interpolation in axial dimension of BIL, DAIN and ZS for actin labeled DCIS cells recorded with a structured illumination microscope. ZS produces better quality results than DAIN and BIL. BIL interpolation results in a faulty elongation of the actin signal highlighted with white arrow. CAFI tools significantly outperform BIL interpolation. Scale bar: 10 µm. Data from Weigert *et al.* (2).

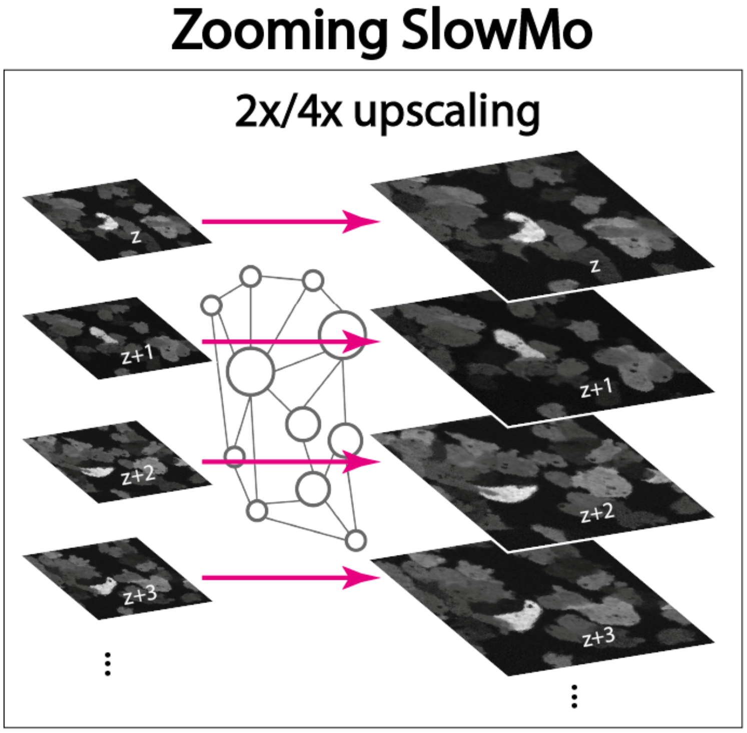


**Fig. S13.** Schematic representation of 2x or 4x lateral upsampling functionality of Zooming SlowMo (ZS). This functionality of ZS was compared for four different datasets by comparing it with BIC, BIL upscaling and two neural network solutions PSSR (1) and SRFBN-S (33).

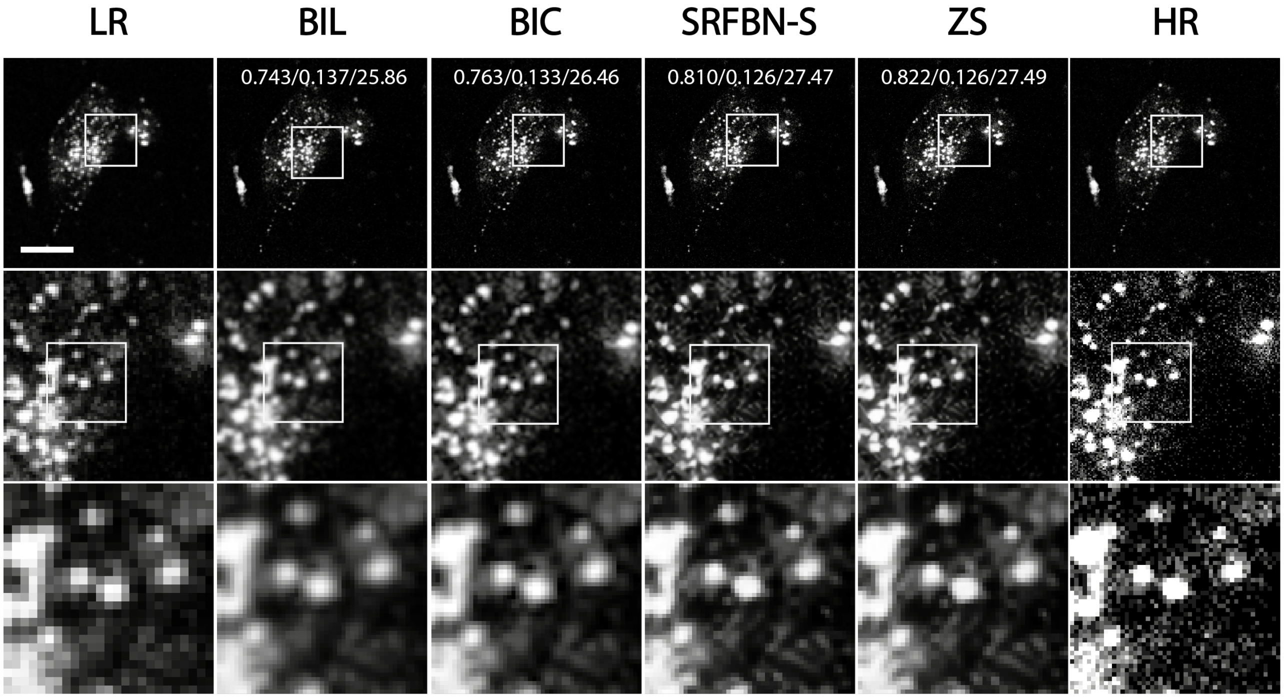


**Fig. S14.** 2x lateral upsampling of lysosomes of SH-SY5Y cells recorded on a confocal microscope (256 px to 512 px). SSIM/RMSE/PSNR values displayed in first row of upsampled images. Scale bar: 15 µm.

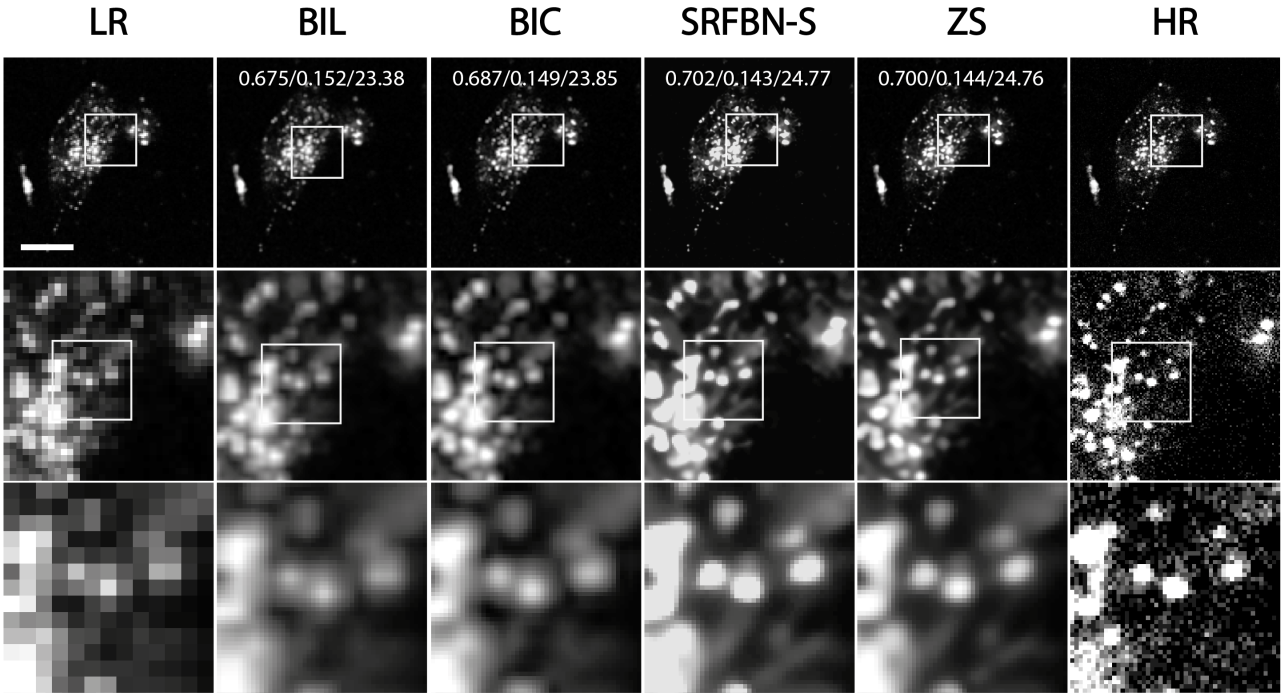


**Fig. S15.** 4x lateral upsampling of lysosomes of SH-SY5Y cells recorded on a confocal microscope (128 px to 512 px). SSIM/RMSE/PSNR values displayed in first row of upsampled images.
Scale bar: 15 µm.

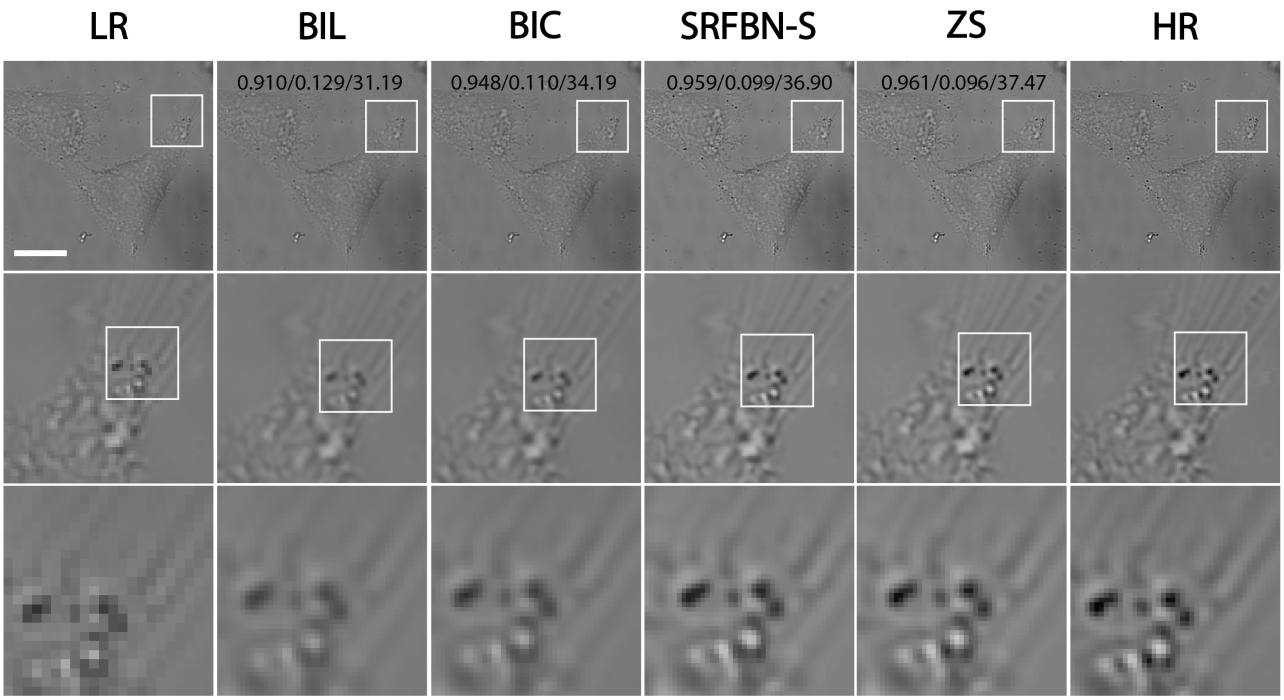


**Fig. S16.** 2x lateral upsampling of SH-SY5Y cells recorded on a confocal brightfield microscope (256 px to 512 px). SSIM/RMSE/PSNR values displayed in first row of upsampled images. Scale bar: 15 µm.

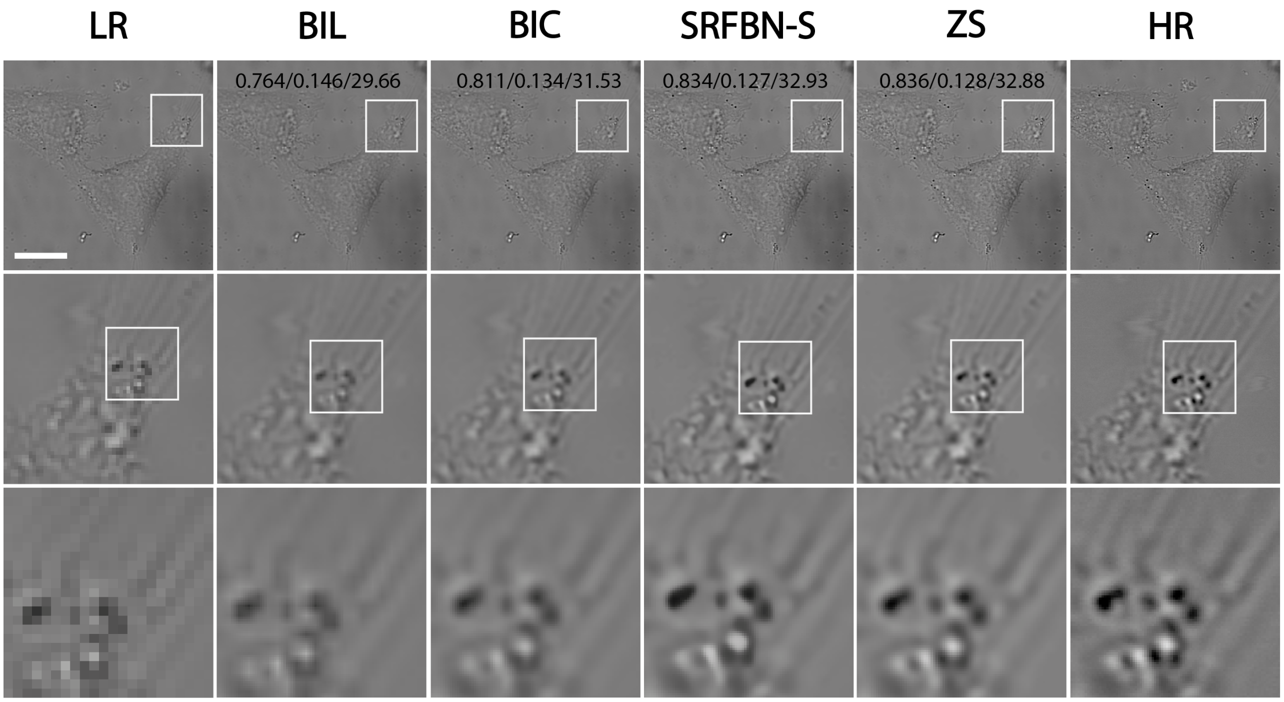


**Fig. S17.** 4x lateral upsampling of SH-SY5Y cells recorded on a confocal brightfield microscope (256 px to 1024 px). SSIM/RMSE/PSNR values displayed in first row of upsampled images. Scale bar: 15 µm.

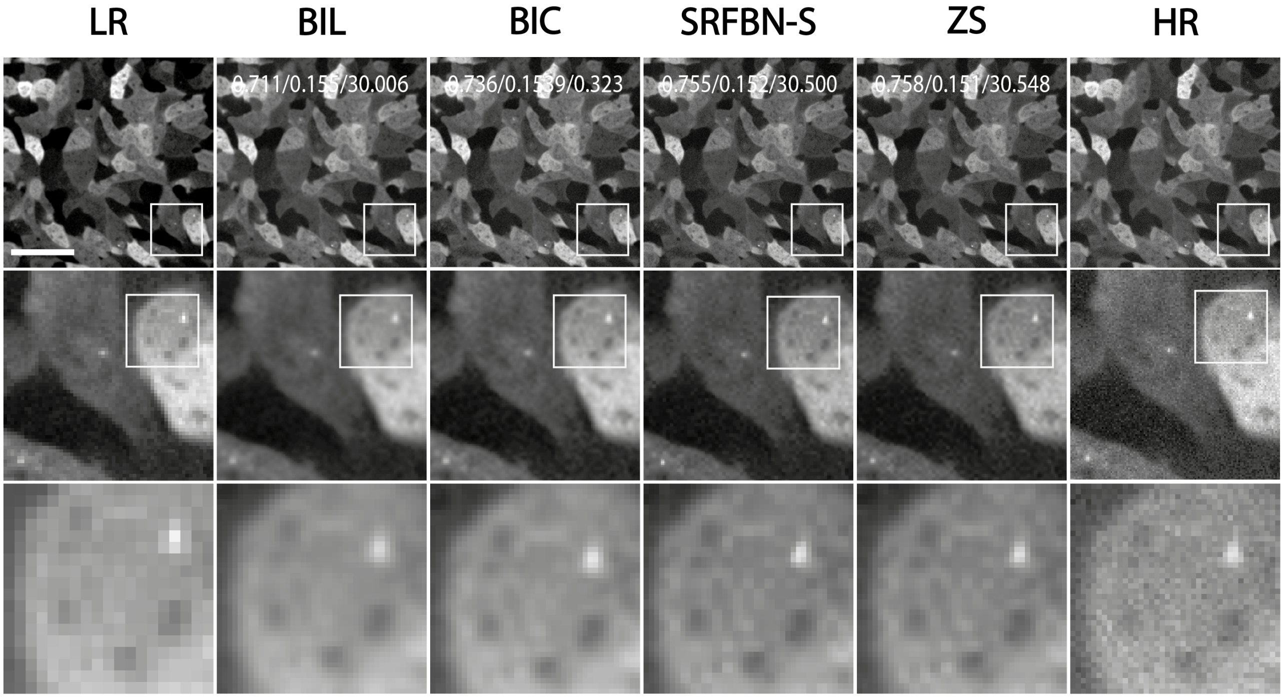


**Fig. S18.** 2x lateral upsampling of dictyostelium recorded on a spinning-disk confocal microscope (256 px to 512 px). SSIM/RMSE/PSNR values displayed in first row of upsampled images. Scale bar: 12 µm.

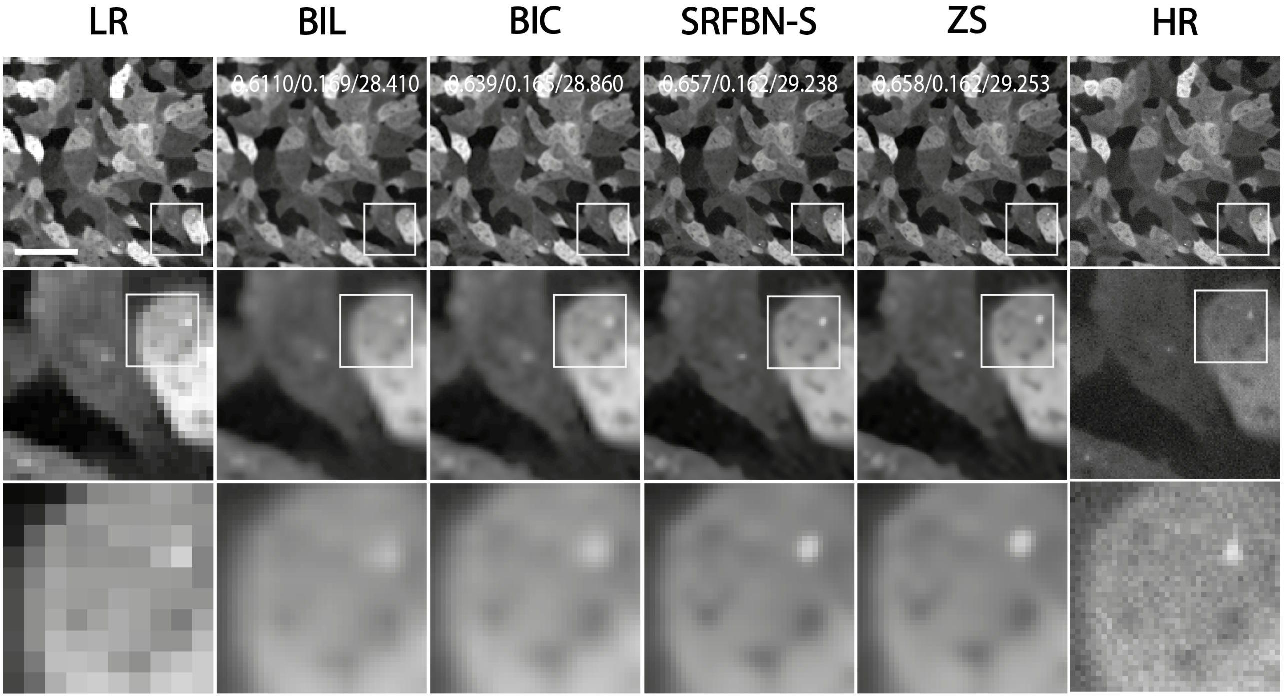


**Fig. S19.** 4x lateral upsampling of dictyostelium cells recorded on a spinning-disk confocal microscope (128 px to 512 px). SSIM/RMSE/PSNR values displayed in first row of upsampled images. Scale bar: 12 µm.

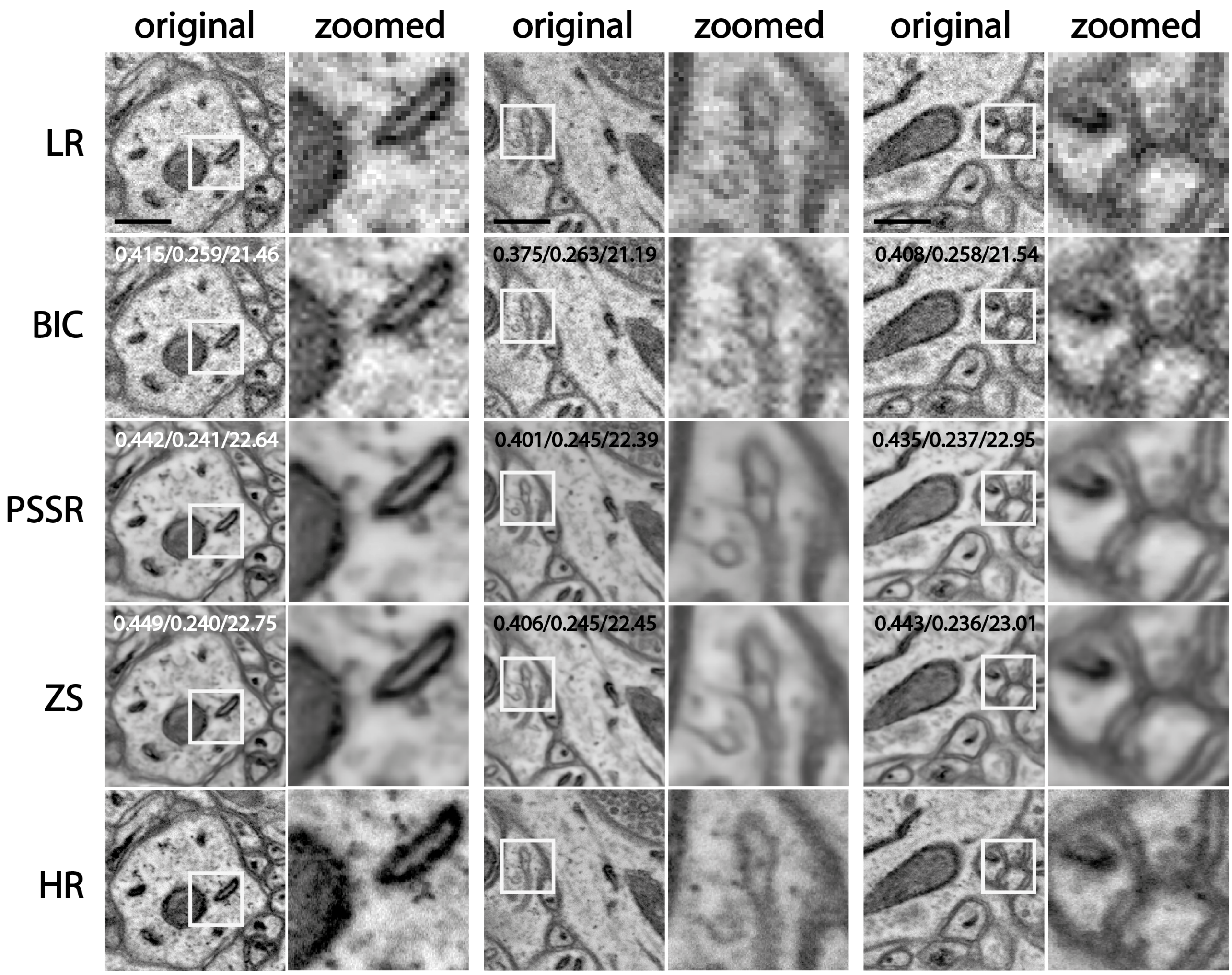


**Fig. S20.** Lateral upsampling comparison. Three representative noisy electron microscopy image examples with 4x lateral upsampling results of BIC, PSSR and ZS compared to low (LR) and high (HR) resolution images. Right image part shows the zoomed in section highlighted with white box in the original image. Quality metrics (SSIM, RMSE, PSNR) are presented in the original lateral upsampling image of each category. Scale bars: 0.4 µm. Data from Fang *et al.* (1).

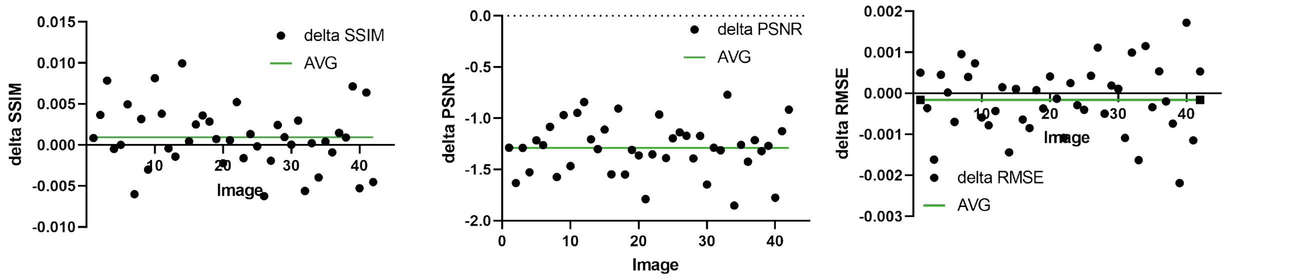


**Fig. S21.** Quality evaluation metrics comparison of all provided 4x lateral upsampling images of PSSR network in comparison to ZS upsampling. Delta SSIM, delta PSNR, and delta RMSE are the differences of the three evaluation metrics by subtracting the evaluated image metric value of PSSR from the metric value of the same image achieved by ZS lateral upsampling.
